# Analysis of pre-symptomatic *Drosophila* models for ALS and SMA reveals convergent impact on functional protein complexes linked to neuro-muscular degeneration

**DOI:** 10.1101/2022.06.20.496821

**Authors:** Marina Garcia-Vaquero, Marjorie Heim, Barbara Flix, Marcelo Pereira, Lucile Palin, Tânia M. Marques, Francisco R. Pinto, Javier de Las Rivas, Aaron Voigt, Florence Besse, Margarida Gama-Carvalho

## Abstract

Spinal Muscular Atrophy (SMA) and Amyotrophic Lateral Sclerosis (ALS) share phenotypic and molecular commonalities, including the fact that they can be caused by mutations in ubiquitous proteins involved in RNA metabolism, namely SMN, TDP-43 and FUS. Although this suggests the existence of common disease mechanisms, there is currently no model to explain the resulting motor neuron dysfunction. In this work we generated a parallel set of *Drosophila* models for adult-onset RNAi and tagged neuronal expression of the fly orthologues of the three human proteins, named Smn, TBPH and Caz, respectively. We profiled nuclear and cytoplasmic bound mRNAs using a RIP-seq approach and characterized the transcriptome of the RNAi models by RNA-seq. To unravel the mechanisms underlying the common functional impact of these proteins on neuronal cells, we devised a computational approach based on the construction of a tissue-specific library of protein functional modules, selected by an overall impact score measuring the estimated extent of perturbation caused by each gene knockdown. Transcriptome analysis revealed that the three proteins do not bind to the same RNA molecules and that only a limited set of functionally unrelated transcripts is commonly affected by their knock-down. However, our integrative approach revealed they exert a concerted effect on protein functional modules, acting through distinct targets. Most strikingly, functional annotation revealed that these modules are involved in critical cellular pathways for motor neurons, including neuromuscular junction function. Furthermore, selected modules were found to be significantly enriched in orthologues of human neuronal disease genes. The results presented here show that SMA and ALS disease-associated genes linked to RNA metabolism functionally converge on neuronal protein complexes, providing a new hypothesis to explain the common motor neuron phenotype. The functional modules identified represent promising biomarkers and therapeutic targets, namely given their alteration in asymptomatic settings.

## Background

Motor neuron diseases (MNDs) are characterized by a progressive and selective degeneration and loss of motor neurons accompanied by an atrophy of innervated muscles. Although MNDs encompass heterogeneous groups of pathologies with different onset and genetic origins, a number of MND-causing mutations have been identified in RNA-associated proteins, leading to a model in which alteration of RNA metabolism may be a key, and potentially common, driver of MND pathogenesis (Achsel et al. 2013; Ling et al. 2013; Taylor et al. 2016; Gama-Carvalho et al. 2017; Zaepfel and Rothstein 2021). This has become particularly clear in the context of two well-studied pathologies: Spinal Muscular Atrophy (SMA) and Amyotrophic Lateral Sclerosis (ALS), which have both been linked to mutations in conserved RNA binding proteins (RBPs). SMA, the most common early-onset degenerative neuromuscular disease, is caused in 95% of patients by a loss of the SMN1 gene, which encodes a protein with chaperone functions essential for the assembly of both nuclear and cytoplasmic ribonucleoprotein (RNP) complexes (Li et al. 2014; Price et al. 2018). The best-characterized role of SMN is to promote the assembly of spliceosomal small nuclear ribonucleoprotein complexes (snRNPs) (Boulisfane et al. 2011; Workman et al. 2012), but it has also been involved in the assembly of other nuclear sRNPs required for 3’end processing (Tisdale et al. 2013), as well as cytoplasmic RNP complexes essential for long-distance mRNA transport (Donlin-Asp et al. 2016; Donlin-Asp et al. 2017). Consistent with these functions, and with additional reported roles in transcription regulation (Pellizzoni et al. 2001; Zou et al. 2004), inactivation of SMN was shown to result in alternative splicing defects (Zhang et al. 2008), transcriptional changes (Zhao et al. 2016) and defective axonal RNA targeting (Fallini et al. 2011; Fallini et al. 2016). To date, how these changes in gene expression account for the full spectrum of symptoms observed in SMA patients and disease models remains unclear. ALS, on the other hand, is the most-common adult-onset MND and has mostly sporadic origins. Remarkably, however, disease-causing mutations in two genes encoding RNA binding proteins, FUS and TDP-43 (alias gene symbol of TARDBP), have been identified in both genetic and sporadic forms of the disease (Da Cruz and Cleveland 2011; Gama-Carvalho et al. 2017). Both proteins shuttle between the nucleus and the cytoplasm and regulate different aspects of RNA metabolism, ranging from transcription and pre-mRNA splicing to mRNA stability and axonal targeting (Ratti and Buratti 2016; Ederle and Dormann 2017; Birsa et al. 2020). ALS-causing mutations were described to have pleiotropic consequences, compromising both the nuclear and cytoplasmic functions of FUS and TDP-43, and resulting in their accumulation into non-functional cytoplasmic inclusions (Ling et al. 2013; Zbinden et al. 2020). Whether ALS pathogenesis primarily originates from a depletion of the nuclear pool of these RBPs, or rather from a toxic effect of cytoplasmic aggregates, has remained unclear (Li et al. 2013; Fernandes et al. 2018).

Thus, SMA and ALS are not only connected by pathogenic commonalities (Bowerman et al. 2018), but also appear to both originate from alterations in RBP-mediated regulatory mechanisms. Further strengthening the possibility that these two MNDs may be molecularly connected, recent studies have suggested that SMN, FUS and TDP-43 belong to common molecular complexes and also exhibit functional interactions (Yamazaki et al. 2012; Groen et al. 2013; Tsuiji et al. 2013; Sun et al. 2015; Perera et al. 2016; Chi et al. 2018; Cacciottolo et al. 2019). Together, these results have raised the hypothesis that SMN, FUS and TDP-43 may control common transcriptional and/or post-transcriptional regulatory steps. The alteration of these common processes in response to an impaired function of either protein would underlie MND progression (Achsel et al. 2013). Comparative transcriptomic studies performed so far, however, did not clearly identify classes of transcripts that may be co-regulated by the three MND RBPs (Lagier-Tourenne et al. 2012; Gama-Carvalho et al. 2017; Kline et al. 2017). This fact has left the question of the existence of common molecular regulatory mechanisms and targets in the diverse MNDs un-answered.

A major difficulty in comparing available transcriptomic studies is that the datasets were obtained from heterogeneous, and often late-stage or post-mortem samples. This prevents robust comparisons and the identification of direct *vs.* indirect targets. Another challenge associated with the identification of relevant regulated mRNAs is that SMN, FUS and TDP-43 are multifunctional and may exhibit distinct sets of target RNAs in the nucleus and the cytoplasm, raising the need for compartment-specific studies. To overcome previous limitations and unambiguously assess the existence of transcripts commonly regulated by SMN, FUS and TDP-43, we decided in this study to systematically identify the direct and indirect neuronal RNA targets of these proteins. For this purpose, we defined a strategy involving the establishment of parallel schemes for tagged-protein expression to perform RNP complex purification, alongside neuron specific, adult-onset gene silencing of the *Drosophila* orthologs of *SMN*, *FUS* and *TARDBP*, identified by the gene symbols *Smn*, *caz* and *TBPH*, respectively.

Highlighting the conservation of protein functions from fly to human, expression of human FUS and TDP-43 proteins was shown to rescue the lethality induced upon inactivation of the corresponding fly genes (Wang et al. 2011). Furthermore, *Drosophila* models based on expression of mutant human or *Drosophila* proteins that recapitulate the hallmarks of SMA and ALS, in particular motor neuron disabilities and degeneration, have been previously established (McGurk et al. 2015; Aquilina and Cauchi 2018; Olesnicky and Wright 2018; Spring et al. 2019; Liguori et al. 2021). Several of these models have been successfully used for large-scale screening and discovery of genetic modifiers (Chang et al. 2008; Kankel et al. 2020; Liguori et al. 2021).

Our study was performed on pre-symptomatic flies starting from head samples. RNA immunoprecipitation sequencing (RIP-seq) experiments were performed to identify the cytoplasmic and nuclear transcripts bound by each protein. These assays were complemented with neuron-specific, adult-onset down-regulation of *Smn*, *caz* or *TBPH* followed by RNA sequencing (RNA-seq) to identify transcripts with altered expression levels and/or splicing patterns. The RIP-seq analysis showed that the three proteins bind to largely distinct sets of RNA targets, in the nucleus and in the cytoplasm. Directly bound mRNAs were not particularly affected by the gene knockdowns, which collectively altered the expression and/or splicing profile of a limited, albeit significant set of common transcripts. These transcripts do not seem to be direct RNA-binding targets and do not present any consistent functional signatures. These observations suggested that the common physiological processes regulated by the three proteins may be altered at a higher order level. To unravel a possible functional relationship between transcripts regulated by Smn, Caz and TBPH, we designed a strategy to map them on neuronal protein complexes. Our results revealed a convergent effect of the three knockdowns on the regulation of common functional modules. Among these modules, we identified seven functional units directly implicated in neuro-muscular junction (NMJ) development. Noteworthy, although these modules were selected by suffering a common impact in all knockdowns, they were also found to be enriched in direct targets identified in RIP-seq experiments. Finally, selected functional modules were enriched in orthologues of human MND-associated genes. In summary, our work provides a new conceptual framework to explain how changes in three ubiquitous proteins involved in RNA metabolism converge into molecular functions critical for MN processes, thereby leading to overlapping disease phenotypes.

## Methods

### Fly lines

The fly stocks used were obtained from the Bloomington *Drosophila* Stock Center (BDSC) and the Vienna *Drosophila* Resource Center (VDRC), or were generated using the *Drosophila* Embryo Injection Service form BestGene (http://www.thebestgene.com). BDSC stocks #39014 (expressing shRNA targeting *TBPH*), #55158 (expressing shRNA targeting *Smn*) and #32990 (expressing shRNA targeting *caz*) were used for the transcriptome profiling assays along with the VDRC strain #13673 (expressing dsRNA targeting *always early*). Transgenic lines used for neuronal expression of GFP-tagged variants of *Smn* (CG16725, fly *SMN*), *TBPH* (CG10327, fly *TARDBP*), *caz* (CG3606, fly *FUS*) were generated by site directed integration into the same attP landing site (VK00013, BDSC#9732).

*Smn*, *caz* and *TBPH* coding sequences were PCR-amplified from ESTs LD23602, UASt-*caz* plasmid (gift from C. Thömmes) and EST GH09868, respectively. Primers used for amplification are listed in Table 1. *Smn* and *caz* PCR products were subcloned into pENTR-D/TOPO vector (Life Technologies), fully sequenced, and recombined into a pUASt-EGFP-attB Gateway destination vector to express N-terminally-tagged proteins. The *TBPH* PCR product was double digested with NotI and XhoI and ligated into a NotI/XhoI digested pUASt-EGFP-attB plasmid (gift from S. Luschnig). Primer sequences are provided in Supplementary Methods.

### Fly crosses

shRNA expression was induced using the GeneSwitch system. Mifepristone was dissolved in 80% ethanol and pipetted on the surface of regular fly food (final concentration of 0,1 mg/cm^2^). Vehicle-only treated fly vials served as control. Vials were prepared 24 hours prior to use to allow evaporation of ethanol. Crosses performed for ‘knockdown’ analyses were as follows: virgins carrying the ubiquitous daughterless GeneSwitch driver (daGS) were crossed with males carrying the UAS:shRNA constructs. In the progeny, male *daGS*/UAS:shRNA flies were collected one day post eclosion (1 dpe) and exposed to food containing mifepristone (replaced every 2nd day). After 10 days, flies were collected, snap frozen in liquid nitrogen and stored at −80°C until further use. Knock-down efficiency of the target genes was assessed by RT-qPCR using rp49 and act5C as normalizing genes using the primers described in Supplementary Methods. The iScript^TM^ cDNA Synthesis Kit (BioRad) was used to transcribe 500ng of total head RNA according to manufacturer’s instructions. Real Time PCR was performed in a total reaction volume of 25 μL using the SYBR^TM^ Green PCR Master Mix (ThermoFisher).

For RIP-seq experiments, males carrying UASt-GFP-fusions (or sole EGFP) were crossed en masse with *elav*-Gal4; tub-Gal80ts virgins. *elav*-Gal4/Y/+; *tub*-Gal80ts/UAS-GFP-*Smn* (or *TBPH* or *caz*) flies were raised at 18°C, switched to 29°C upon eclosion and aged for 5 to 7 days before being collected in 50 mL Falcon tubes and snap frozen.

### Immuno-histochemistry and Western-blotting

For analysis of GFP-fusion distribution, brains were dissected in PBS and immuno-stained using anti-GFP antibodies (1:1,000; Molecular Probes, A-11122), as described previously (Vijayakumar et al. 2019). Samples were imaged on an inverted Zeiss LSM710 confocal microscope. For analysis of GFP-fusion expression, heads were smashed into RIPA buffer (15 heads for 100 μL RIPA) and lysates directly supplemented with SDS loading buffer (without any centrifugation). Total protein extracts or RIP extracts were subjected to SDS Page electrophoresis, blotted to PVDF membranes, and probed with the following primary antibodies: rabbit anti-GFP (1:2,500; #TP-401; Torey Pines); mouse anti-Tubulin (1:5,000; DM1A clone; Sigma); and mouse anti-Lamin (1:2,000; ADL 67.10 and ADL 84.12 clones; DHSB).

### RNA Immunoprecipitation assays

Falcon tubes with frozen flies were chilled in liquid nitrogen and extensively vortexed to separate heads, legs and wings from the main body. Head fractions were collected at 4°C, through sieving on 630 µm and 400µm sieves stacked on top of each other. 1 mL of heads was used per condition, except for GFP-Smn, where 2 mL of heads were used. For the GFP control, 500 µL of heads were mixed with 500 µL of w^1118^ heads to normalize the amount of GFP proteins present in the initial lysate.

Adult *Drosophila* heads were ground into powder with liquid nitrogen pre-chilled mortars and pestles. The powder was then transferred to a prechilled 15 mL glass Dounce Tissue Grinder and homogenized in 8.5 mL of Lysis buffer (20mM Hepes pH 8, 125mM KCl, 4mM MgCl2, 0.05% NP40, 1mM dithoithreitol (DTT), 1:100 Halt™ Protease & Phosphatase Inhibitor Cocktail, Thermo Scientific, 1:200 RNasOUT™, Invitrogen). Cuticle debris were eliminated by two consecutive centrifugations at 100 g for 5 minutes at 4°C. Nuclear and cytoplasmic fractions were then separated by centrifugation at 900 g for 10 minutes at 4°C. The supernatant (cytoplasmic fraction) was further cleared by two consecutive centrifugations at 16,000 g for 20 minutes. The pellet (nuclear fraction) was washed with 1 mL of Sucrose buffer (20 mM Tris pH 7.65, 60 mM NaCl, 15 mM KCl, 0.34 M Sucrose, 1 mM dithoithreitol (DTT), 1:100 Halt™ Protease & Phosphatase Inhibitor Cocktail, Thermo Scientific, 1:200 RNasOUT™, Invitrogen), centrifuged at 900 g for 10 minutes at 4°C and resuspended in 2 mL of Sucrose buffer. 800 µL of High salt buffer (20 mM Tris pH 7.65, 0.2 mM EDTA, 25% Glycerol, 900 mM NaCl, 1.5 mM MgCl2, 1 mM dithoithreitol (DTT), 1:100 Halt™ Protease & Phosphatase Inhibitor Cocktail, Thermo Scientific, 1:200 RNasOUT™, Invitrogen) were then added to reach a final concentration of 300 mM NaCl. After 30 minutes incubation on ice, the nuclear fraction was supplemented with 4.7 mL of Sucrose buffer to reach a concentration of 150 mM NaCl and with CaCl_2_ to reach a 1 mM CaCl_2_ concentration. RNAse free DNase I (Ambion™, Invitrogen) was added (0.1 mM final concentration) and the sample was incubated for 15 minutes at 37°C with gentle agitation. EDTA was added to a 4 mM final concentration to stop the reaction, and the digested fraction was centrifuged at 16,000 g for 20 minutes (4°C) to obtain soluble (supernatant; used for immuno-precipitation) and insoluble (pellet) fractions.

Cytoplasmic and nuclear fractions were incubated for 30 minutes at 4°C under agitation with 120 µL of control agarose beads (ChromoTek, Germany). Pre-cleared lysates were collected by a centrifuging 2 min at 400g (4°C). Immuno-precipitations (IPs) were performed by addition of 120 µL of GFP-Trap®_A beads (ChromoTek, Germany) to each fraction and incubation on a rotator for 1.5 hours at 4°C. Tubes were then centrifuged for 2 minutes at 2,000 rpm (4°C) and the unbound fractions (supernatants) collected. Beads were washed 5 times with Lysis buffer, resuspended in 100 µL of Lysis buffer supplemented with 30 µg of proteinase K (Ambion) and incubated at 55°C for 30 minutes. Eluates (bound fractions) were then recovered and further processed. At least three independent IPs were performed for each condition.

### RNA extraction, Library preparation and RNA sequencing

RNA from IP eluates or frozen fly heads (50 flies aprox/genotype) was extracted using Trizol according to the manufacturer’s instructions. RIP-Seq libraries were prepared in parallel and sequenced at the EMBL Genomics core facility. Briefly, libraries were prepared using the non-strand-specific poly(A)+ RNA Smart-Seq2 protocol (Nextera XT part). Following quality control, cDNA libraries were multiplexed and sequenced through single-end 50 bp sequencing (HiSeq 2000, Illumina).

For the RNA-preparation used in RNA-Seq experiments, 30-50 fly heads were homogenized in 0.5 ml Trizol (ThermoFischer) using a speed mill (Analytic Jena) and ceramic beads. Debris was removed by short centrifugation and supernatant was transfered to a fresh Eppendorf tube and incubated at 10 minutes on ice. 0.2 ml Chloroform were added and mixture was vortexed for 15 seconds and incubated on ice for 10 minutes followed by centrifugation for 15 minutates, 17.000 xg at 4°C. Upper phase was removed and volume determined. RNA was precipitated by addition of 1x volume Isopropanol and incubated on ice for 1 hour follwed by a centrifugation for ten minutes with 17.000 xg at 4°C. Supernatant was removed and pellet was washed twice with 1 ml ice cold 70% ethanol, air dried and resuspended in 25 µl RNAse free water.

RNA-seq libraries for shRNA analysis were prepared and sequenced at the Genomics Facility, Interdisziplinäres Zentrum für Klinische Forschung (IZKF), RWTH Aachen, Germany. Libraries were generated using the Illumina TrueSeqHT library protocol and ran on a NextSeq machine with paired-end sequencing and a read length of 2×76nt. The 47 raw fastq files of the RNA-seq data generated for this study have been submitted to the European Nucleotide Archive under the umbrella project FlySMALS, with accession numbers PRJEB42797 and PRJEB42798.

### RNA-seq data analysis

Following quality assessment using FastQC version 0.11.5 (https://www.bioinformatics.babraham.ac.uk/projects/fastqc/), all raw sequencing data was processed with in-house perl scripts to filter out reads with unknown nucleotides, homopolymers with length ≥50 nt or an average Phred score < 30, and trim the first 10 nucleotides (Amaral et al. 2014) Remaining reads were aligned to the BDGP *D. melanogaster* Release 6 genome assembly build (dos Santos et al. 2015) using the STAR aligner version 2.5.0 (Dobin et al. 2013) with the following options: –outFilterType BySJout –alignSJoverhangMin 8 –alignSJDBoverhangMin 5 –alignIntronMax 100000 –outSAMtype BAM SortedByCoordinate –twopassMode Basic – outFilterScoreMinOverLread 0 –outFilterMatchNminOverLread 0 –outFilterMatchNmin 0 – outFilterMultimapNmax 1 –limitBAMsortRAM 10000000000 –quantMode GeneCounts. Gene counts were determined using the htseq-count function from HTseq (version 0.9.1) in union mode and discarding low quality score alignments (–a 10), using the Flybase R6.19 annotation of gene models for genome assembly BDGP6.

For RIP-seq data analysis, gene counts were normalized and tested for DE using the DESeq2 (Love et al. 2014) package of the Bioconductor project (Huber et al. 2015), following removal of genes with less than 10 counts. mRNAs associated with each protein were identified by performing a differential expression analysis (DEA) for each condition vs the corresponding control pull-down. Transcripts with a positive log2 FC and an adjusted p value for DEA lower that 0.05 were considered to be bound by the target protein.

DEA for RNA-Seq gene counts was performed with the limma Bioconductor package (Ritchie et al. 2015) using the voom method (Law et al. 2014) to convert the read-counts to log2-cpm, with associated weights, for linear modelling. The design formula (∼ hormone + Cond, where hormone = treated or non-treated and Cond = Caz, Smn or TBPH RNAi) was used to consider hormone treatment as a batch effect. Differential gene expression analysis was performed by comparing RNAi samples for each target protein to control (always early RNAi) samples. Genes showing up or down-regulation with an adjusted p value <0.05 were considered to be differentially expressed.

Altered splicing analysis (ASA) was performed on the RNA-seq aligned data using rMATS version 4.0.2 (Shen et al. 2014) with flags -t paired --nthread 10 --readLength 66 --libType fr-firststrand. For the purpose of the downstream analysis, the union of all genes showing any kind of altered splicing using the junction count and exon count (JCEC) analysis with a FDR <0.05 in the comparison between each target gene RNAi versus control RNAi was compiled as a single dataset.

Normalized RNA-Seq data of adult fly brain tissue was retrieved from FlyAtlas2 database in November 2020 (www.flyatlas2.org; (Leader et al. 2018)). Neuronal transcripts were filtered applying an expression threshold of >1 FPKM (Fragments Per Kilobase per Million). This gene set was then used to filter the final gene lists from RIP-seq, DEA and ASA. The full universe of 8,921 neuronal genes is annotated in Supplementary Data 5.

Clustering analysis was performed using the heatmap function from ggplot2 R package (default parameters) (Whickam 2016) and correlation plots were generated using lattice R package. Intersection analyses of RNA-Seq and RIP-seq datasets were performed using UpSetR and SuperExactTest R packages (Wang et al. 2015).

### Network analysis and generation of the library of functional modules

*Drosophila* physical Protein-Protein Interaction (PPI) data was retrieved from APID repository (http://apid.dep.usal.es; (Alonso-López et al. 2016, Alonso-López et al. 2019)) in December 2019. The original unspecific network was filtered to include only interactions between proteins expressed in adult fly brain tissue as described in previous section. The neuronal network was then simplified to remove self-loops and isolated proteins using the igraph R package (Mora and Donaldson 2011). Bioconductor GOfuncR R package was used to evaluate the functional enrichment of brain network as compared to the unspecific network - Gene Ontology Biological Process, hyper-geometric test, FDR = 0.1 on 1000 randomizations-(Grote 2020). Finally, the functional modules were defined by selecting groups of physically interacting proteins annotated under the same enriched term. It should be noted that not all the proteins collaborating in the same process must physically interact (*e.g.,* as in the case of cell signaling, the membrane receptor does not bind to its downstream transcription factor). Based on this, we enabled modules to be formed by non-connected subnetworks. The isolated clusters were discarded only when the largest subnetwork represented more than 90% of the total module. The same protein might be annotated with several terms and therefore might be involved in several modules simultaneously. Conversely, we are aware that the use of GO data may return functionally redundant modules. Prior any further analysis, module redundancy was evaluated to check that modules do not exceedingly overlap nor represent redundant biological processes. Based on this analysis, a module size from 10 up to 100 proteins was defined as optimal to minimize redundancy.

## Results

### Caz, Smn and TBPH proteins do not share common mRNA targets

We hypothesized that the existence of shared RNA targets for Caz, Smn and TBPH might underlie the observed phenotypic commonalities between SMA and ALS. To test this hypothesis, we performed RIP-seq to identify neuronal mRNAs present in the RNP complexes formed by each of these proteins in adult *Drosophila* neurons. To facilitate cross-comparisons and ensure reproducible and cell-type specific purification, we generated three independent transgenic lines with GFP-tagged constructs expressed under the control of UAS sequences inserted into the same chromosomal position. To specifically characterize the neuronal RNA interactome, GFP-fusion proteins were expressed in adult neuronal cells using the pan-neuronal *elav*-GAL4 driver. The ectopic expression of Caz, Smn and TBPH has been reported to induce toxicity (Grice and Liu 2011; Xia et al. 2012; Cragnaz et al. 2014). For this reason, we used the TARGET method (McGuire et al. 2003) to express GFP-fusion proteins specifically in adult neurons within a limited time window (5-7 days post-eclosion). The TARGET system relies on the temperature-sensitive GAL80 protein, which inhibits GAL4 at low temperature, enabling temporal regulation of UAS constructs. When expressed in neuronal cells, GFP-Caz and GFP-TBPH robustly accumulated in the soma, showing a predominant, although not exclusive nuclear accumulation (Supplementary Fig. S1A and S1C). As expected, GFP-Smn was found mainly in the cytoplasm, sometimes accumulating in foci (Supplementary Fig. S1B). Despite the same insertion site and promotor sequence, GFP-Smn protein was consistently expressed at lower levels (Supplementary Fig. S1D).

Since Caz, Smn and TBPH are multifunctional proteins involved in both nuclear and cytoplasmic regulatory functions, we separately characterized their RNA interactome in each cellular compartment. For this purpose, cellular fractionations were performed prior to independent anti-GFP immunoprecipitations, thus generating paired nuclear and cytoplasmic samples (Fig. 1A). As shown in Fig. 1B, relatively pure nuclear and cytoplasmic fractions were obtained from head lysates and GFP-tagged proteins could be efficiently immuno-precipitated from each fraction.

**Figure 1.**
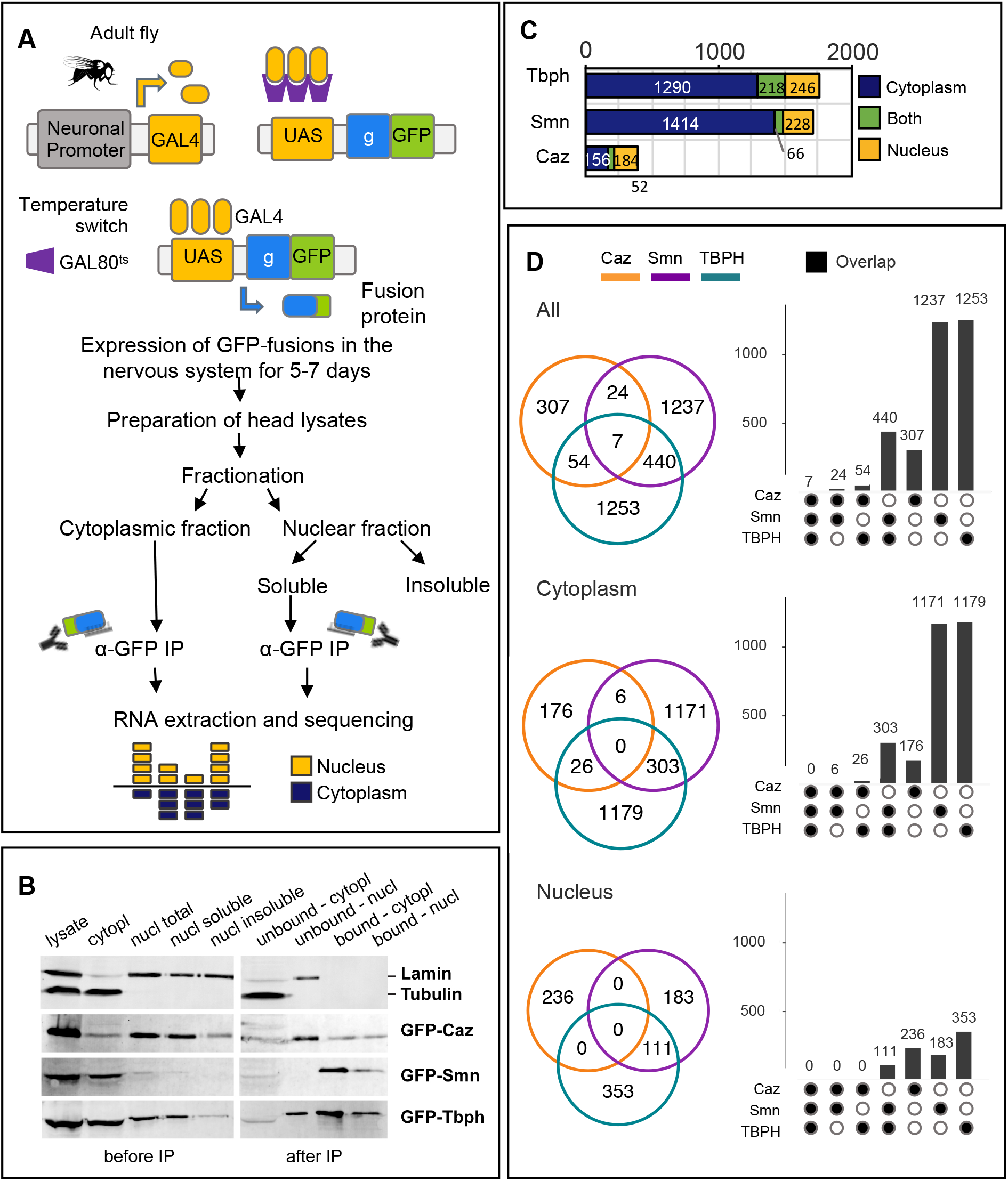
RIP-Seq identification of mRNAs bound by Caz, Smn or TBPH in adult *Drosophila* neurons. (A) Schematic representation of the RIP-Seq procedure. GFP-fusion proteins were conditionally expressed in the nervous system of adult flies *via* the Gal4/Gal80/∼UAS system. Head lysates were prepared and fractionated into cytoplasmic and nuclear fractions (see Methods). Nuclear proteins were further solubilized with high salt buffer and recovered in the soluble fraction. The cytoplasmic and nuclear soluble fractions were used for immuno-precipitation with GFP-trap beads. Co-immunoprecipitated mRNAs were extracted and sequenced. (B) Western blot analysis of the different fractions obtained in the RIP procedure. Lamin and Tubulin were used as markers of the nuclear and cytoplasmic fractions, respectively. Depletion of Tubulin from the nuclear fraction and of Laminin from the cytoplasmic fraction demonstrates the quality of the fractionation procedure. Also note the differential distribution of GFP-Caz, Smn and TBPH in the different fractions. As expected, Smn was predominantly localized in the cytoplasmic fraction, and Caz in the nuclear. In contrast, TBPH was abundant in both fractions. (C) Bar graph showing the number of neuronal mRNAs co-immunoprecipitated with Caz, Smn or TBPH from cytoplasmic (blue) or nuclear (yellow) lysates. Transcripts found in both compartments are shown in green. (D) Venn diagrams and corresponding bar graphs illustrate the overlap between the total (top panel), cytoplasmic (middle panel) or nuclear (lower panel) mRNA interactomes of Caz, Smn and TBPH. The seven transcripts found in the overlap of the top diagram (“All”) correspond to mRNA molecules present in distinct compartments, in agreement with no common transcripts being found in the cytoplasmic and nuclear fraction overlap analysis.

For each paired nuclear and cytoplasmic pull-down, co-precipitated RNAs were extracted and used to prepare mRNA-seq libraries. Pull-downs from flies expressing GFP were used as control. Three independent replicate datasets were generated for each protein, except for GFP-Caz, for which one nuclear pull-down sample did not pass quality control for library generation. The raw sequencing dataset, composed of 23 libraries containing between 17.7 and 64.6 million total reads (Supplementary Data 1), was submitted to the European Nucleotide Archive (ENA) with the study accession code PRJB42798.

RIP-seq datasets were analyzed to identify mRNA molecules enriched in GFP-fusion versus GFP control pull-downs with an average of 13,500 genes detected across all samples, ranging (Supplementary Data 1). Sequencing datasets clustered primarily depending on the nuclear *versus* cytoplasmic natures of the extract, and secondly depending on the protein used for pull-down (Supplementary Fig. S2A). Differential expression analysis (DEA) was performed between each of the six pull-down conditions and the corresponding GFP control, with transcripts displaying positive enrichment (adjusted p value < 0.05) considered as associating with the target protein (Supplementary Data 2).

Although Caz, Smn and TBPH fusion proteins were expressed specifically in neurons *via* the *elav* promotor, a certain degree of RNP complex re-association may occur in head lysates during the different experimental steps, as previously described (Mili and Steitz 2004). To discard any non-neuronal transcripts that may have co-precipitated with target proteins, the dataset resulting from the DEA was filtered to include only genes with reported expression in the adult fly brain (see Methods), corresponding on average to 70% of the enriched transcripts (see Supplementary Fig. S3).

These analyses revealed that Smn and TBPH associate with a large fraction of the neuronal transcriptome (1,708 and 1,754 mRNAs in total, respectively), and that most of their identified mRNA targets associate in the cytoplasm rather than in the nucleus (Fig. 1C). A much smaller number (208) of mRNAs were found to associate with Caz in the cytoplasm, with 236 mRNAs detected as enriched in the pull-downs from nuclear fractions. Although this may partly reflect the higher heterogeneity of the Caz pull-down samples (Supplementary Fig. 2A), it is in good agreement with the low abundance of GFP-Caz protein found in the cytoplasm compared to GFP-Smn and GFP-TBPH (Fig. 1B). Of note, the percentage of transcripts simultaneously bound by the same protein on both compartments averaged only 22%, with TBPH displaying a much larger overlap than Smn for a similarly sized set of target mRNAs (Fig. 1C). This observation is in agreement with the current model of mRNP complex remodeling between the nucleus and the cytoplasm, with the compartment-specific set of mRNA bound proteins being influenced both by their relative affinities and abundance (Mili and Steitz 2004).

We next addressed the existence of common RNA targets, which could provide insights regarding a potential common MN degenerative mechanism associated with the altered expression of the human orthologue proteins in a disease context. Overlap analysis of the mRNA interactomes of Caz, Smn, and TBPH revealed a striking absence of transcripts bound by all three RBPs in the cytoplasmic or nuclear fractions (Fig. 1D). This finding might not exclusively result from the small number of RNAs bound by Caz, as a poor overlap was also observed between the large sets of cytoplasmic mRNAs bound by TBPH and Smn. Considering that the universe of protein-associated transcripts was defined exclusively based on the adjusted p value, without imposing a minimal enrichment threshold, this observation is particularly surprising. Together, our RIP-seq experiments thus uncovered that Caz, Smn and TBPH do not share common RNA targets.

### Gene expression changes in response to reduced levels of Caz, Smn and TBPH have significant commonalities but lack a clear functional signature

In addition to regulatory roles associated with mRNA binding activity, Caz, Smn, and TBPH have been shown to have both direct and indirect roles as transcriptional, translational, and splicing regulators (reviewed in Gama-Carvalho 2017). It is thus possible that, despite associating to non-overlapping sets of mRNAs, these proteins may coordinate common gene expression programs through other molecular mechanisms. To address this hypothesis, we used shRNA-expressing fly lines to knock-down the expression of *caz*, *Smn*, and *TBPH* in adult flies by RNA interference (RNAi) and characterized the resulting changes in neuronal gene expression using RNA-seq (Fig. 2A). After identification of fly lines displaying a robust silencing of each target gene, we used the GeneSwitch (GS) system to induce ubiquitous, adult-onset RNAi (Osterwalder et al. 2001). This system relies on the feeding of flies with the hormone mifepristone (RU486), which activates GAL4-progesterone-receptor fusions, thus driving expression of the shRNA transgenes (Fig. 2A). Given that the system has been reported to display some leakage in the absence of the hormone (Scialo et al. 2016), a fly line expressing shRNA against the non-related embryonic transcript *always early* (ae) was used as control. Three to five days post-eclosion, resulting male progeny was transferred to food with or without the shRNA-inducing hormone for ten days and knock-down efficiency of target genes was evaluated by qRT-PCR (Supplementary Fig. 1E). Of note, despite an effective knock-down of the target gene levels of ∼50%, these flies did not exhibit any motor phenotype or increased mortality. Therefore, our model represents a pre-symptomatic stage of the molecular pathways regulated by the *caz*, *Smn* and *TBPH* genes. Total RNA extracted from fly heads from three independent experimental assays was used for library preparation and paired-end Illumina mRNA sequencing (RNA-seq). The raw sequencing dataset, composed of 24 libraries with an average number of 50 million reads (Supplementary Data 1), was submitted to ENA with the study accession code PRJB42797.

**Figure 2.**
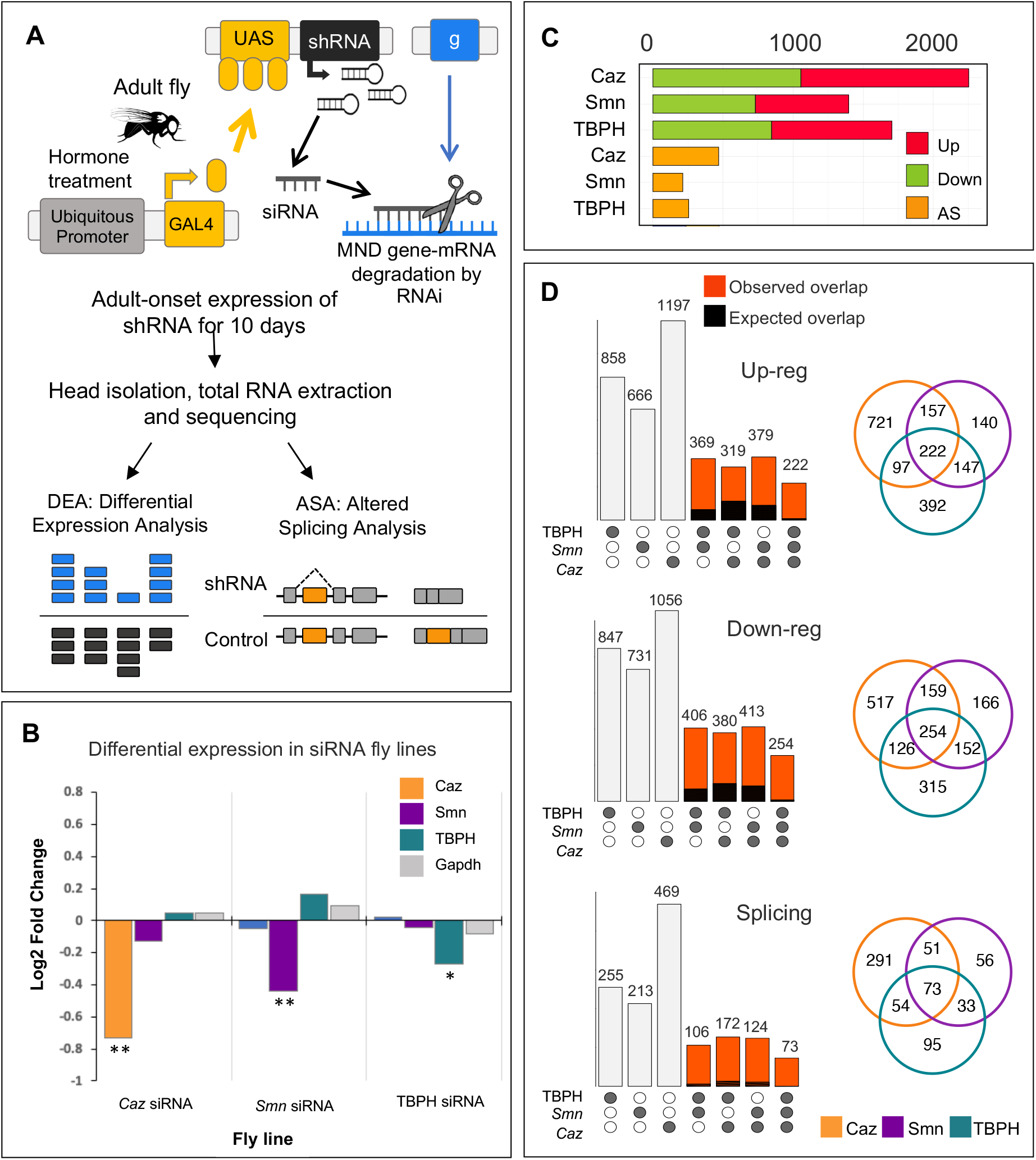
Identification of differentially expressed neuronal transcripts in response to RNAi-mediated silencing of Caz, Smn or TBPH in adult flies. (A) Schematic representation of the experimental set-up for RNA-seq analysis. Hormone-dependent, adult-onset expression of short hairpin (sh) RNA was used to induce RNAi-mediated gene silencing of *caz*, *Smn* or *TBPH*. RNA was prepared from fly heads five to seven days post induction of shRNA expression, quality checked and subjected to mRNA-seq. Fly lines with shRNA against the *always early* (ae) embryonic gene served as control. (B) Bar graph showing the RNA-seq log2 fold change of the siRNA target genes plus the Gapdh housekeeping gene in each RNAi fly line. Statistically significant differences to the *ae* siRNA fly line are indicated as ** (adjusted p value < 0.05) and * (adjusted p value = 0.06 and p value < 0.02). (C) Bar graph showing the number of upregulated (red), downregulated (green) and differentially spliced mRNAs (yellow) in each RNA-seq dataset. Note that the kind of splicing change was not considered for this analysis. (D) Venn diagrams and corresponding bar graphs showing the overlap in upregulated (top), downregulated (middle) or differentially spliced (bottom) mRNAs in flies with RNAi-mediated silencing of *caz*, *Smn* or *TBPH.* The bar color compares the expected (black) and observed (red) overlap given the total transcripts altered in response to the silencing of *caz*, *Smn* or *TBPH,* respectively. The expected ratios were calculated using *SuperExactTest* R package.

Following quality control and filtering, reads were aligned to the *Drosophila* reference genome, mapping to ∼13,600 expressed genes. The RNA-seq dataset was analyzed to determine the overall changes in transcript abundance and splicing patterns induced by the knock-down of each protein. Exploratory analysis of the normalized RNA-seq dataset revealed that the samples clustered primarily according to genotype, followed by treatment (Supplementary Fig. S2B), an observation consistent with the expected leakage from the siRNA locus. However, hormone-treated samples exhibited a better separation between genotypes than the corresponding untreated controls, as expected from shRNA-expressing samples (Supplementary Fig. S2C,). As hormone treatment induced a significant number of common changes across all sample types, explaining up to 7% of the variance in the dataset (Supplementary Fig. S2C), differential gene expression (DE) analysis was performed between hormone treated *Caz*, *Smn* and *TBPH* shRNA-expressing target and control fly lines (see Supplementary Data 3). Confirming the robustness of our dataset and DE analysis, the specific shRNA target genes were found to be significantly down-regulated exclusively in the corresponding fly line (Fig. 2B). Given that our three target proteins are known to regulate mRNA processing, we also analyzed the data to identify alternative splicing (AS) changes that occurred as a consequence of the gene knockdowns (Supplementary Data 4). Although the analysis we performed distinguishes between five distinct types of AS, for the aim of the present study all AS changes identified in each siRNA line were combined and transcripts defined as either alternatively spliced, or not affected. Taking into consideration that RNA-seq was performed using samples isolated from fly heads, the list of transcripts showing significant DE or AS changes in response to *caz*, *Smn* or *TBPH* knock-down was filtered as previously described to exclude non-neuronal genes (Supplementary Fig. S3). Supplementary Data 5 provides the final annotated list of all neuronal genes detected in the different fly models and experiments.

Fig. 2C summarizes the overall results of the RNA-seq analysis. Although the proportion of up- and down-regulated genes within the DE gene set (∼50%) was similar in all conditions, there were considerable differences in the number of DE or AS transcripts identified in response to each knockdown (Fig. 2C). Given the correlation between these numbers and the observed knock-down efficiency and sample heterogeneity (Fig. 2B and Supplementary Fig. S2B), we believe these differences more likely reflect our experimental set-up than a specific characteristic of the gene expression programs regulated by each target protein. Importantly, only a relatively small fraction of the DE transcripts was identified as a direct protein target in the RIP-seq assays (Supplementary Data 5). Caz-regulated transcripts showed minimal direct association with Caz protein (4.6%), whereas ∼22% of the genes showing altered expression in response to Smn or TBPH RNAi were found to be enriched in the corresponding RIP-seq assays. Interestingly, this fraction goes up to ∼40% when considering only the transcripts displaying AS changes in response to Smn or TBPH knock-down, suggesting the splicing of these transcripts may be directly regulated by each protein (Supplementary Data 5).

We next looked for the commonalities between the sets of transcripts with altered expression induced by the knockdown of each target protein. A summary of the number of genes displaying common changes in expression as a consequence of the shRNA knockdowns is depicted in Fig. 2D. The overlap analysis of these gene sets revealed that approximately 500 genes exhibit similar gene expression changes in response to all knockdowns (Fig. 2D). This is well above the overlap expected by random chance, with an estimated p value < 1e-16. Performing the overlap analysis for common DE genes without requiring that the type of change is identical in the three fly lines only retrieved an additional 20 genes, underscoring the significance of the common changes that were detected. Thus, despite the total lack of common RIP-seq targets, the downregulation of Caz, Smn and TBPH protein expression seems to elicit a coherent transcriptome response.

In an attempt to understand the functional consequences of this common transcriptome signature, we performed a functional enrichment analysis to identify altered biological processes. Surprisingly, almost no Gene Ontology (GO) terms were enriched in the subset of ∼500 common genes (Supplementary Data 6). This result is in stark contrast with the strong functional signature that was observed for GO enrichment analysis of the subsets of mRNAs captured in the RIP-Seq assays. Furthermore, the identification of functional signatures was not improved by imposing more stringent fold change cut-offs on the DE gene sets. Indeed, the majority of DE genes showing common changes across all knockdowns presents very small variations in expression (Supplementary Fig. 2D). The use of more stringent fold change cut-offs on our dataset leads to a big reduction in the size of the gene sets (Supplementary Fig. 2E), with an even larger impact on the number of common DE genes that are retained (Fig. 2F). These observations support our choice to consider all genes showing statistically significant changes in expression in our analysis, independently of their fold change level. However, we were unable to link this group of genes to a coherent set of molecular processes that are directly influenced by the three proteins, which might provide novel insights into the human disease context. Nevertheless, given the clear presence of a shared transcriptome signature, we hypothesized that putative underlying functional connections might be obscured by independent, albeit converging phenomena, taking place at the protein network level. To obtain insights into these potential connections, we proceeded to a more in-depth network-based analysis of our datasets.

### Network-based approaches identify commonly affected neuronal functional modules

Biological processes are dynamic and complex phenomena that emerge from the interaction of numerous proteins collaborating to carry out specialized tasks. Thus, a biological process can be similarly impacted by changes in distinct proteins that contribute to the same regulatory function.

To understand whether the phenotypic commonalities observed in ALS and SMA might result from the deregulation of distinct, but functionally connected target proteins, we used a computational network-based approach. First, following our recently published approach (Garcia-Vaquero et al. 2022), we generated a library of tissue-specific “functional modules” comprised of physically interacting and functionally collaborating neuronal proteins (Fig. 3A). To do so, we began by reconstructing the entire *Drosophila* neuronal interaction network using protein-protein interaction (PPI) and adult fly brain RNA-seq datasets available in the APID and FlyAtlas2 repositories, respectively (Leader et al. 2018; Alonso-López et al. 2019). Notably, 45.5% of the 5 353 proteins found in this neuronal network are encoded by transcripts whose levels and/or splicing were altered in response to *caz*, *Smn* and/or *TBPH* knockdowns. Next, we defined functional modules in the neuronal network by selecting groups of physically interacting proteins annotated under the same enriched functional term. Of the 232 modules with associated GO terms, we focused on the subset of 122 modules composed of 10 to 100 proteins (Supplementary data 7). These modules retained 1541 proteins in total, maintaining the high percentage of Caz, Smn and/or TBPH-dependent genes found in the original network (43.7%).

**Figure 3.**
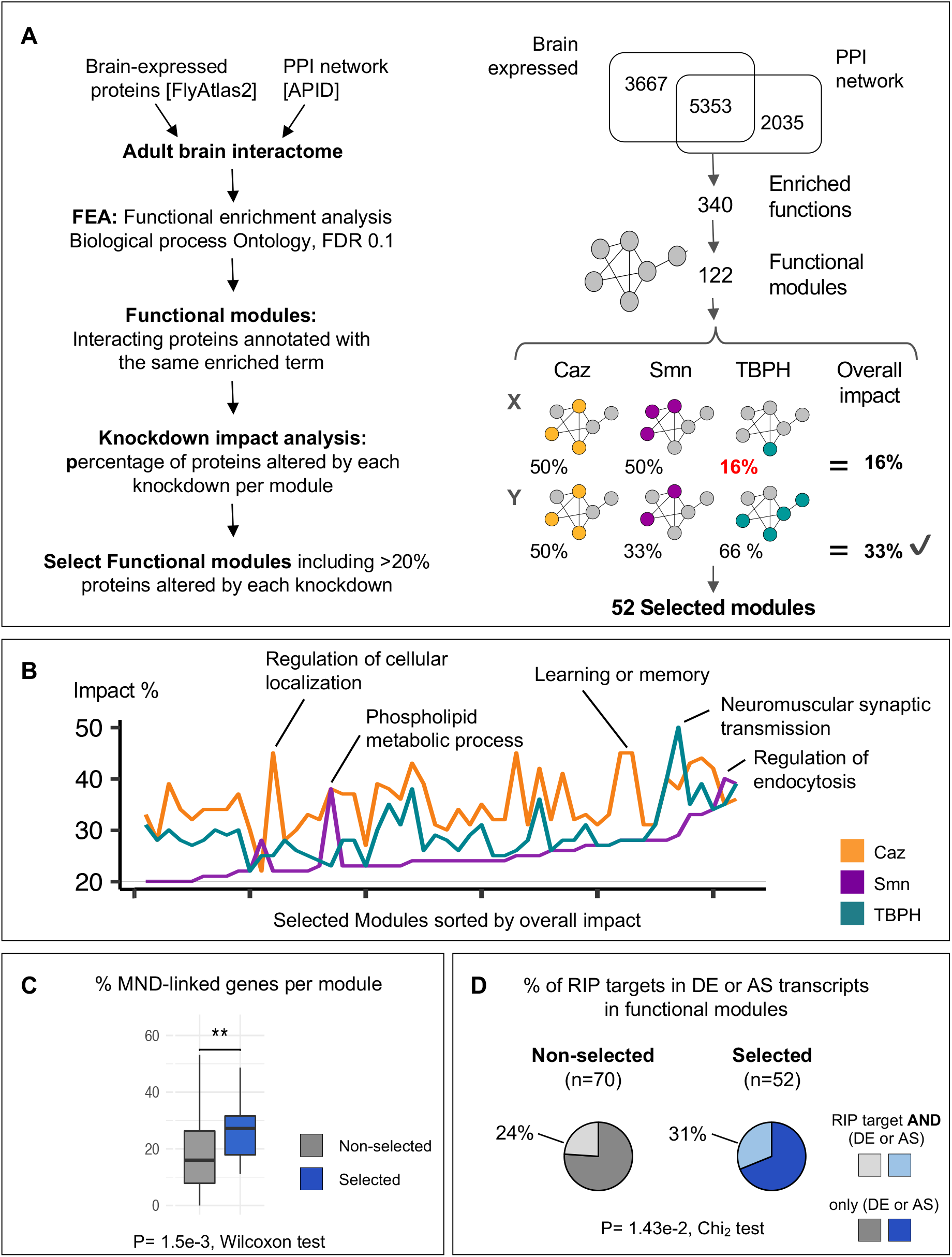
Characterization of functional modules impacted by reduced abundance of Caz, Smn and TBPH proteins. (A) Left panel: workflow used to generate and select functional modules. The adult brain interactome was obtained from the APID protein-interaction network after filtering for proteins expressed in the adult *Drosophila* brain (i, see Methods). Functional enrichment analysis of the resulting interactome was performed to retrieve overrepresented GO Biological Processes (ii). Note that the functional enrichment returns all the proteins annotated in each overrepresented term. The modules were generated from the functional enrichment by retaining the proteins annotated and simultaneously interacting in the brain network (iii). Finally, the impact of *caz, Smn* and *TBPH* knockdown was evaluated for each module (iv) to select modules with > 20% of transcripts altered in each individual knockdown (v). Right panel: summary the workflow outputs. “Overall impact” calculation is exemplified for two modules (X/Y) with the impact score indicated on the right. Only the Y module would be selected, as the overall impact of module X is below the defined threshold. (B) Line plot comparing the impact of individual knockdowns on selected modules, sorted by increasing overall impact. Modules with the highest impact for each protein are indicated by their short name. (C) Box plots showing the percentage of proteins with MND-linked orthologs in each module class. Selected modules (blue) are significantly enriched in proteins with MND-linked orthologs compared to non-selected modules (grey) (p value = 1.5e-3, Wilcoxon test). (D) Pie charts representing the fraction of transcripts with altered expression (DE) or splicing (AS) in response to a given protein knockdown that are simultaneously found in RNP complexes bound by the same protein. The high percentage of DE/AS transcripts (selected modules, blue pie chart) is significantly related to a higher frequency of DE/AS transcripts involved in RBPs bound by the same proteins (p-value = 5.4e-3, Chi^2^ test of independence).

To evaluate the impact of each of the three proteins on individual functional modules, we calculated the percentage of nodes belonging to the DE or AS categories. To focus on modules simultaneously affected by the downregulation of *caz*, *Smn* and *TBPH*, we assigned to each module an “overall impact” score, defined as the minimal percentage of transcripts showing altered expression in any given knockdown (Fig. 3A). 52 modules with an overall impact score of ≥ 20% were identified. These modules were selected for further analysis, as they seem to be under the common control of all three proteins, although not necessarily through regulation of the same target genes.

Consistent with the potential functional relevance of the selected modules, associated functional terms were found to comprise a range of biological processes relevant in a MND context. These include general cellular processes such as kinase signal transduction pathways, regulation of the actin cytoskeleton, regulation of endocytosis, as well as neuron-specific processes such as learning and memory, and regulation of synapse assembly (Supplementary Data 7). Interestingly, differences in the impact of individual gene knockdowns were observed when comparing modules, which we propose to reflect some degree of functional specialization of the two ALS-related genes and the single SMA-associated gene (Fig. 3B). For example, the module related to “learning and memory” functions was strongly impacted by *caz* down-regulation, but to a lower extent by *Smn* or *TBPH* silencing. In contrast, the module “neuromuscular synaptic transmission” was strongly impacted by *TBPH*, followed by *caz*, and less so by *Smn* knockdown. Finally, some modules, like the one linked to “regulation of endocytosis” tended to be similarly impacted by all three knockdowns. Overall, the impact of *TBPH* and *Caz* knockdowns on functional modules is much more similar to each other than to *Smn*, which generally displays lower impact scores, with a few exceptions including “regulation of endocytosis” (Fig. 3B). This observation is quite striking considering that *Caz* and *TBPH* are associated to the same disease.

To determine the relevance of the selected modules to the pathophysiology of MNDs, we calculated for each module the percentage of proteins with human orthologs already linked to MNDs (according to the DisGeNET repository). Remarkably, selected modules were significantly enriched in proteins with MND-linked human orthologs when compared to modules that did not pass the “overall impact” threshold (*p* value = 1.5e-3, Wilcoxon test) (Fig. 3C). This result suggests that we were able to identify novel disease-relevant interactions based on the convergent analysis of *Caz*, *Smn* and *TBPH*-dependent functional modules in *Drosophila*.

As the selected modules represent core biological functions regulated by the three proteins, we looked at the prevalence of direct targets (i.e mRNAs identified by RIP-seq) among module components classified as DE and/or AS in the knockdown analysis. We found that 31% of the 411 DE/AS transcripts associated to selected modules are also bound by at least one of the three MND proteins. In non-selected modules, this number decreases significantly to 24% (*p* value = 1.4e-2, Fig. 3D), being even lower for transcripts that do not integrate any functional module (18% of 2280 transcripts, *p* value = 3.1e-8; Supplementary Data 7). Together, these results suggest that our integrated data analysis approach was able to identify key functional processes that are commonly and directly regulated by the three proteins. Our findings point to the ability of Caz, Smn and TBPH protein dysregulation to elicit a convergent functional impact through distinct individual targets. The connection to key biological processes is mediated by functional protein networks enriched in molecules with known links to MNDs. Further exploration of the selected networks may thus provide novel information to understand MND pathophysiology.

### Convergent disruption of neuromuscular junction processes by altered Caz, TBPH or Smn protein levels

Pairwise comparison of the 52 selected modules revealed a high number of shared genes between many of them (see Supplementary Fig. S4). To generate a non-redundant map of the common functional networks established by Caz, Smn and TBPH, we coalesced groups of highly interconnected modules into larger but more condensed “super-modules” (Fig. 4). This resulted in seven super-modules named after their core functional association: signaling, traffic, cytoskeleton, stress, behavior, synaptic transmission, and neuro-muscular junction (NMJ) (Supplementary Data 7). These super-modules range in size from 77 to 259 nodes, with a maximum overlap between any two super-modules of ∼12% of the nodes (Supplementary Fig. S4). We next determined the presence of MND-associated gene orthologues in the different super-modules (MND-linked, Fig. 5, left panel). We further mapped the distribution of DE transcripts that are direct targets of Caz, Smn and TBPH (RNA-binding, Fig. 5, middle panel); and of transcripts showing altered splicing (Altered Splicing, Fig. 5, right panel). This analysis revealed a distinctive distribution of these characteristics in the groups of modules that were coalesced into super-modules, which is particularly evident regarding the percentage of transcripts displaying altered splicing or with potential roles in MND. The super-modules related to behavior, neuro-muscular junction (NMJ) and cytoskeleton incorporated the largest fraction of MND-linked and AS transcripts. Given the critical link between MNDs and the physiology of NMJs, we focused on the NMJ super-module for a more in-depth analysis.

**Figure 4:**
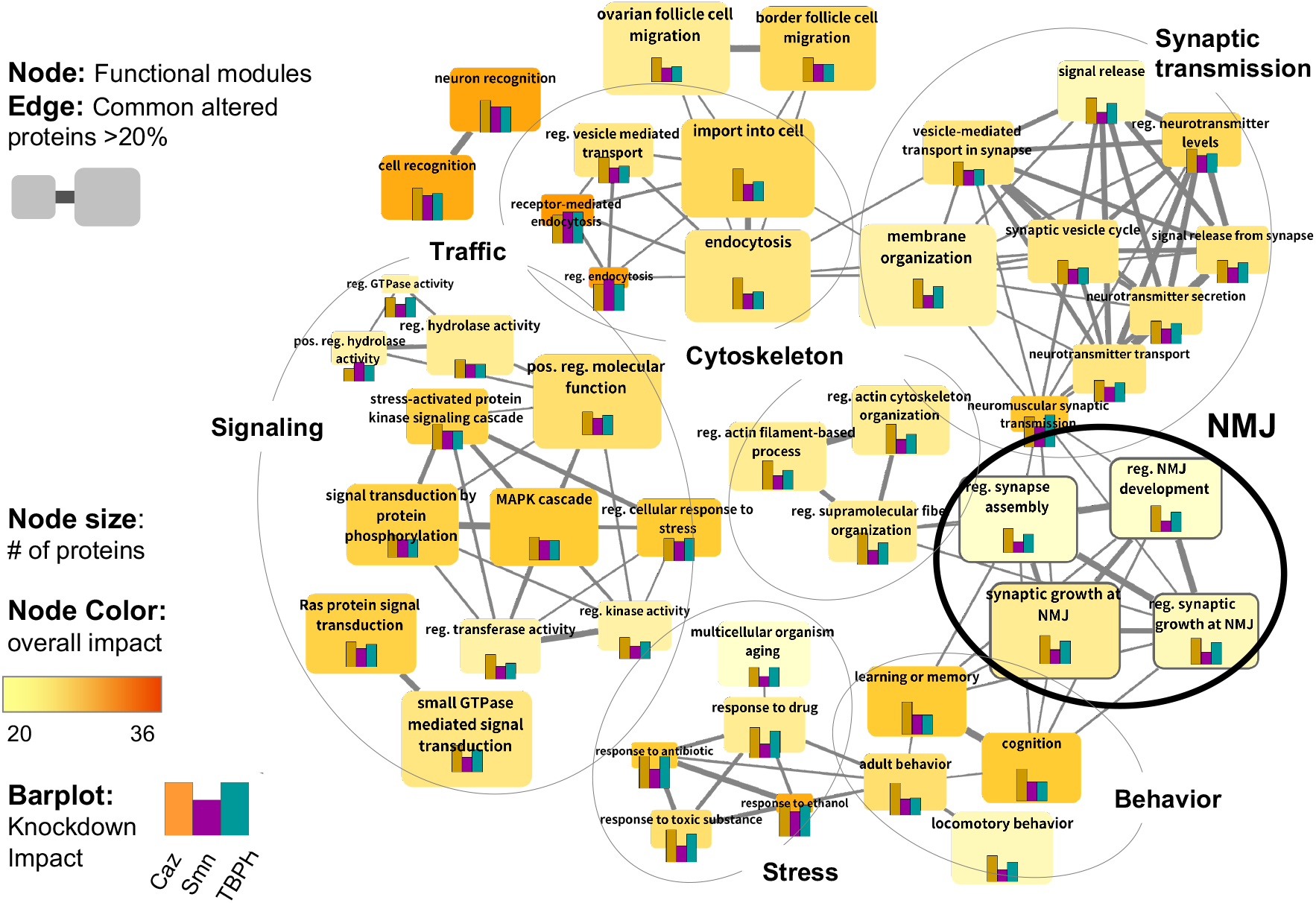
Identification of functional super-modules through protein overlap analysis. (A) Network representation of the selected functional modules. Nodes represent the selected modules designated by the original name of the gene ontology term. Node size indicates the number of proteins incorporated in the module and gradient color the overall impact, i.e., minimum % of transcripts altered by each knockdown. The bar plots within the nodes indicate the impact of each knockdown on the module. Edge width indicates the number of commonly altered transcripts between two modules. Modules were manually grouped into 7 “super-modules” (circles) based on edge density (common altered transcripts) and functional similarity of module names.

**Figure 5:**
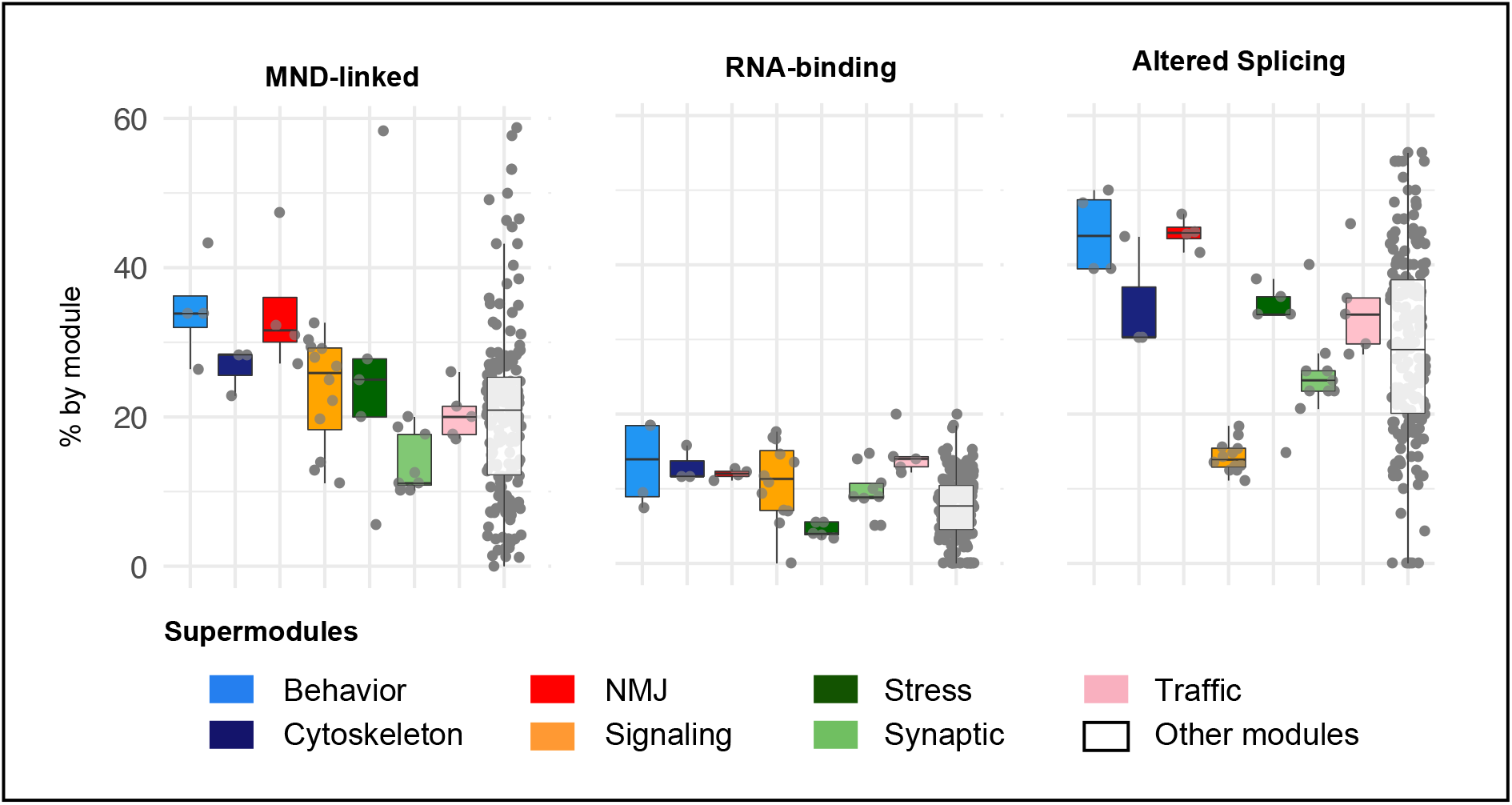
Analysis of super-module features. Box plots showing the distribution of the percentage of annotated proteins in the different super-modules (colored boxes) and across all other (non-selected) library modules (white boxes). Grey dots represent the same percentage in the individual modules that are part of the super-module group. (Left) Percentage of proteins encoded by MND-linked gene orthologs according to the DisGeNET repository. (Middle) Percentage of proteins encoded by DE or AS transcripts that are direct RNA-binding targets of Caz, Smn or TBPH. (Right) Percentage of proteins encoded by transcripts with altered splicing patterns.

The NMJ super-module comprises 104 proteins, of which 49% (51 nodes) are encoded by genes differentially expressed and/or displaying altered splicing in at least one knockdown condition (Supplementary Data 8). 38 of these genes establish direct interactions, forming the subnetwork represented in Fig. 6A. To assess the degree to which the NMJ “super module” functionally interacts with Caz, Smn and TBPH *in vivo*, we cross-referenced it to genetic modifiers of *Drosophila* Smn, Caz or TBPH mutants identified in genome-wide screens for modulators of degenerative phenotypes using the Exelixis transposon collection (Kankel et al. 2020, Sen et al 2013, Chang et al 2008). Interestingly, 21 nodes (∼20%) of the NMJ “super module” were identified as either suppressors or enhancers of these models of neurodegeneration (Supplementary Data 8). Given that the reported percentage of recovered modifiers in these screens ranged between 2% and 5%, this result highlights the biological relevance of the functional modules identified through our approach.

**Figure 6:**
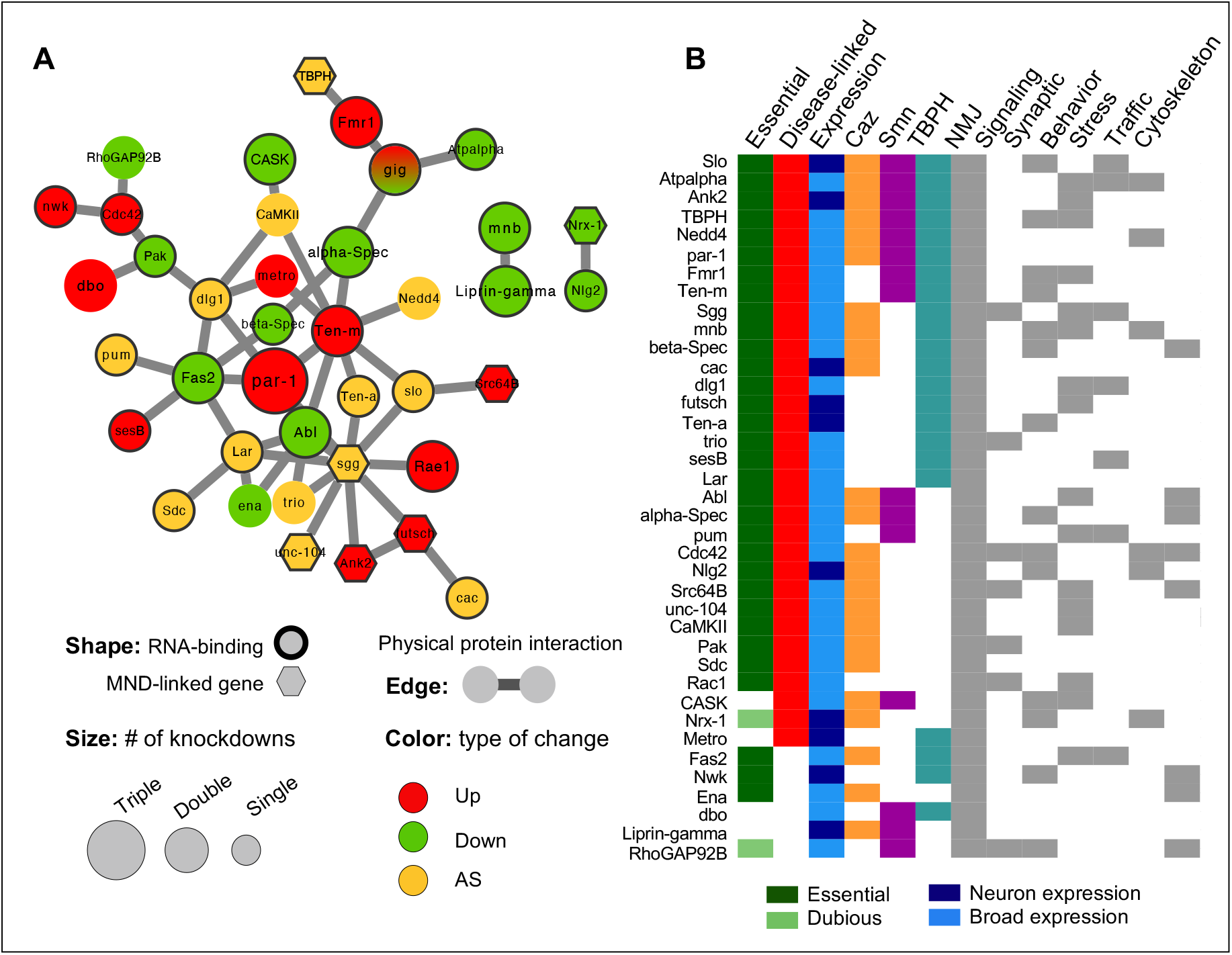
The neuro-muscular junction (NMJ) super-module. (A) Protein-interaction subnetwork of NMJ super-module nodes that are encoded by transcripts altered by knockdown of Caz, Smn and/or TBPH. Only proteins with direct interaction with other proteins encoded by DE/AS transcripts are represented. Node size indicates the number of knockdown models in which the transcript revealed altered expression (DE) and/or splicing (AS). Several transcripts are both DE and AS; yellow nodes indicate transcripts only showing AS. Bold outline highlights proteins encoded by transcripts identified in the RIP-Seq analysis. Hexagons highlight proteins with MND-linked human orthologs. (B) Categorical heat map summarizing data concerning the proteins/genes within the network represented in A. Essential proteins were defined according to the FlyBase repository. Proteins labeled as “Dubious” display a lethal phenotype after induction of RNAi. Thus, it is likely that flies homozygous for amorphic mutations would result in lethality during development. However, since this might result from off-target effects, they were not considered essential. MND-associations were retrieved from the DisGeNET repository. Caz, Smn and TBPH columns indicate in which knockdown models the corresponding transcripts were found altered. Last 7 columns indicate whether the protein is also found in other super-modules.

Detailed analysis of the FlyBase annotations for the genes within the NMJ subnetwork represented in Fig. 6A provides interesting insights into the potential mechanisms causing neuronal dysfunction in the context of MNDs.

First, essential genes are highly overrepresented in the module. While about 30% of *Drosophila* genes are expected to be essential for adult viability (Spradling et al. 1999), more than 75% of genes present in the NMJ super-module have a lethal phenotype (Fig. 6B). Exceptions are *CASK*, *liprin-γ*, *Nlg2*, *metro*, *dbo* and *nwk*. For *RhoGAP92B* and *Nrx-1*, it is so far not entirely clear whether mutant alleles would cause lethality.

We next asked whether the human orthologs of these genes are linked to neurological disorders. *TBPH* (TDP-43), *unc-104* (KIF1A, B, C), *Ank2* (Ank2), *futsch* (MAP1A/B), *sgg* (GSK3A/B), *Src64B* (FYN/SRC) and *Nrx-1* (Nrx-1-3) have been implicated in MNDs (hexagonal nodes in network). Moreover, a high number of genes have human orthologs linked to other neuronal dysfunctions or diseases. For example, human orthologs to fly genes *CASK* (CASK), *Mnb* (DYRK1A), *Rac1* (RAC1), *Dlg-1* (DLG1), *Cdc42* (CDC42), *Fmr1* (FMR1, FXR1/2), *trio* (TRIO), *Nedd4* (NEDD4L/NEDD4) and *CamKII* (CAMK2A/B/D) have been linked to intellectual disability. Epilepsy has been associated with mutations in the human gene orthologs of *cac* (CACNA1A/B/E), *alpha-Spec* (SPTAN1) and *slo* (KCNMA1). In addition, human psychiatric diseases like schizophrenia or bipolar disorder can be caused by alterations in genes with high similarity to *Pak* (PAK1/2/3) and *dbo* (KLHL20 indirect, via regulation of Pak, (Wang et al. 2016)). Alterations in the human gene coding Teneurin Transmembrane Protein 4 (TENM4, shares high homology with fly *Ten-a* and *Ten-m*) are known to cause hereditary essential tremor-5, while human neuroligins NLGN1, NLGN3 and NLGN4X were linked to autism/Asperger syndrome and encode orthologs to fly *Nlg2*. Finally, alterations in human orthologs to fly *Pum* (PUM1/2), *beta-Spec* (SPTBN1/2) and Ank2 (ANK1/2/3) have been associated with Ataxia-like phenotypes and mental retardation. In total, we were able to find direct associations to human MN or neurological disorders for 32 out of the 38 represented genes. Thus, although most of the genes captured in our analysis are not exclusively expressed in neurons, their mutations are somehow associated to abnormal neuroanatomy and function. Interestingly, this holds true for the non-essential genes as well. It is also noteworthy that, in spite of the relatively limited overlap between the different super-modules, all the proteins that constitute this core NMJ network are common to at least another super-module, and on average to more than half of them (Fig 6B).

Altogether, these observations imply that the proteins encoded by the NMJ super-module genes fulfill relevant functions in NMJ maintenance and that their alteration could eventually contribute to MND. Our results reveal that *Caz*, *Smn* and *TBPH* act in concert to regulate biological processes linked to NMJ maturation and function by altering the expression of transcripts encoding distinct, yet physically and functionally interacting proteins. We propose that the functional complexes established by these proteins may represent important players in disease progression, emerging as potential common therapeutic targets rather than the individual proteins that compose them.

## Discussion

SMA and ALS are the most common MNDs and are characterized by a progressive degeneration of motor neurons and loss of skeletal muscle innervation. Although both diseases share many pathological features, including selective motor neuron vulnerability, altered neuronal excitability, as well as pre- and post-synaptic NMJ defects (Bowerman et al. 2018), their very different genetic origins and onset led them to be classified as independent, non-related diseases. This view has been challenged by recent studies demonstrating that disease-causing proteins (Smn for SMA, Fus and TDP-43 for ALS) are connected through both molecular and genetic interactions (reviewed by Gama-Carvalho et al, 2017). Furthermore, the increasing number of functions attributed to these proteins converges onto common regulatory processes, among which control of transcription and splicing in the nucleus, as well as mRNA stability and subcellular localization in the cytoplasm. Despite the observed convergence in the molecular function of Smn, Fus and TDP-43, transcripts co-regulated by these three proteins, and thus central to SMA and ALS pathophysiology, have not been identified by previous transcriptomic analyses. In this study, we used the power of *Drosophila* to systematically identify, on one hand the mRNA repertoires bound by each protein in the nucleus and cytoplasm of adult neurons and, on the other hand, the mRNA populations undergoing significant alterations in steady-state levels or splicing as a consequence of the knockdown of each protein. This approach revealed a striking absence of mRNAs commonly bound by the three proteins and a small, albeit significant, number of commonly altered transcripts. Notwithstanding, and contrary to the simplest model that explains shared disease phenotypes, this subset of shared transcripts did not present any functional signature linking it to biological pathways related to disease progression.

Considering that functional protein complexes are at the core of all critical cellular mechanisms, an alternative model posits that shared phenotypes may arise through convergent effects on independent elements of such complexes. To investigate this possibility, we mapped the de-regulated transcripts identified in our transcriptomic analysis onto a comprehensive and non-biased library of neuronal physically interacting and functionally collaborating protein consortia. Following our recently published approach (Garcia-Vaquero 2022), definition of these functional units was achieved through the integration publicly available information from *Drosophila* PPI networks, neuronal gene expression and gene ontology annotations. This approach led to the identification of a set of 52 functional modules significantly impacted by all three proteins through the regulation of distinct components (Fig. 3). Of note, although we used as selection criterium the presence of a minimum of 20% of module elements displaying altered gene expression in each knock-down model, we found that modules passing this cut-off were significantly enriched in direct RNA binding targets of Smn, Caz and TBPH compared to non-selected modules (Fig. 3D). Considering that only a very small proportion of these targets are common to the three proteins, this observation underscores our hypothesis of convergent regulation of functional complexes through distinct individual elements. Furthermore, the enrichment of RIP targets in the selected modules establishes a direct mechanistic link between changes in the levels of Smn, Caz and TBPH and changes in the steady state expression of module components. It is possible that the steady-state levels of transcripts encoding other proteins that belong to the same complex will vary as part of homeostatic feed-back processes. This could justify the presence of a relatively large number of DE/AS genes that are common to the three knockdown models, but whose transcripts are not found as direct protein targets in our RIP-seq data.

The functional classification of the 52 selected modules revealed a striking connection with critical pathways for MND. Particularly relevant, mapping of the human orthologues of the different module components revealed a high number of genes with reported association to MNDs. This observation provides support to the relevance of our approach, which uses *Drosophila* as a model for uncovering molecular interactions underlying human disease. It is noteworthy that the enrichment in disease-associated orthologues was not homogeneous across the super-modules generated by coalescing highly related modules into a smaller number of larger functional protein consortia (Fig. 5). Interestingly, we found that a super-module related to NMJ function was among the highest scoring regarding both enrichment in MND associated genes and presence of alternatively spliced/direct RNA binding targets. The subset of DE/AS genes present in this module forms a highly interconnected network and the analysis of FlyBase annotations for this focused subset provided interesting insights into potential mechanisms that may underlie neuronal disfunction. An unusually large number of DE/AS genes within the NMJ super-module was found to correspond to essential genes, indispensable for the development of adult flies. Alterations in the abundance and/or function of these genes have been linked in several cases to a disturbance of nervous system function. This is reflected by an alteration in stress response and/or abnormal behavior in either embryos, larvae or adult flies. Strikingly, even the non-lethal genes captured in this super-module have been shown to impact nervous system development and cause abnormal neuroanatomy when mutated/silenced.

The essential function of most of the selected genes obviously prohibits the analysis of loss-of-function phenotypes in the adult organism. In neurons, classical forward and reverse genetics of essential genes and clonal analysis is complex. This is the reason why there is little genetic data on gene products involved in neuronal maintenance. Conditional knockouts and spatiotemporal control of RNAi-mediated gene silencing (like the approach used here) is a way to overcome these limitations. We can only speculate whether a neuron specific, adult-onset knockdown of the individual genes within the super-module will impair adult neuron integrity. However, it is reasonable to assume that the collective deregulation of this set of genes within the super-module is incompatible with proper neuronal function. This assumption is particularly sound if the encoded proteins and their associated functional complexes are found to contribute to cellular processes critical for neurons, as indeed we find in this case. In fact, for almost all proteins encoded by the NMJ sub-network, synaptic functions have been reported. Interestingly, the other identified super-modules are also functionally annotated to cellular mechanisms that are especially important in neurons, like signaling, cytoskeletal dynamics, traffic and transport. Thus, an attractive model emerges for SMA and ALS MN dysfunction: convergent functional impacts can emerge from the independent, subtle deregulation of a group of proteins that are part of a set connected, neuronal functional modules. A persisting impairment in critical neuronal processes could initiate a self-reinforcing cycle of detrimental events, eventually resulting in neuronal decline. Especially in the case of sporadic, late-onset ALS, this model would comply with the events observed in disease progression.

In conclusion, our work revealed common functional modules that are under the control of the SMA and ALS disease-associated gene orthologues *Smn*, *TBPH* and *caz*. This control is exerted through distinct target genes that encode proteins which collaborate in neuronal functional consortia. The fact that these modules are deregulated in pre-symptomatic disease models and are primarily composed of ubiquitously expressed genes opens an interesting avenue of research regarding the discovery of novel disease biomarkers. Importantly, the identification of convergent molecular dysfunctions linked to distinct MND-associated genes suggests that common therapeutic strategies able to help slowdown disease progression or improve symptoms may exist, in spite of the diversity of genetic backgrounds.

## List of Abbreviations

ALS: Amyotrophic Lateral Sclerosis
AS: Alternative Splicing
DE: Differentially Expressed
FC: Fold Change
GFP: Green Fluorescent Protein
GO: Gene Ontology
MN: Motor Neuron
MND: Motor Neuron Disease
NMJ: Neuromuscular Junction
RBP: RNA Binding Protein
RIP: RNA Immuno-Precipitation
RNAi: RNA interference
RNA-seq: RNA sequencing
RNP: Ribonucleoprotein
PPI: Protein-Protein-Interaction
shRNA: short hairpin RNA
SMA: Spinal Muscular Atrophy

## Declarations

### Ethical Approval and Consent to participate

Not applicable

### Consent for publication

Not applicable

### Availability of supporting data

The datasets generated and/or analyzed during the current study are available in the European Nucleotide Archive repository under the umbrella study FlySMALS, with accession numbers PRJEB42797 (https://www.ebi.ac.uk/ena/browser/view/PRJEB42797) and PRJEB42798 (https://www.ebi.ac.uk/ena/browser/view/PRJEB42798).

## Acknowledgements

The authors would like to acknowledge Jörg B. Schulz and Joachim Weiss for support and helpful discussions during the course of the project. We thank the Genomics Core Facility (EMBL, Germany) for assistance with library preparation from RIP samples and sequencing.

## Authors’ contributions

MG-C, AV, FB, and JDLR conceptualized the research approach and supervised the research work; BF validated and generated the RNAi samples; MH generated and characterized the GFP-tagged lines and did the RNA-IP assays; LP assessed the functionality of GFP-tagged lines; MP and TMM did the RNA-seq data analysis; MG-V developed the methods and performed the network-based analysis; FRP contributed to the conceptualization and development of the network analysis. MG-C, AV, FB and MG-V wrote the manuscript draft. All authors read and approved the final manuscript.

## Authors’ Information

Not applicable

## Additional files

### Additional file 1

**Supplementary Figure S1. Characterization of the UAS-GFP-Caz, UAS-GFP-Smn and UAS-GFP-TBPH fly lines.** (A-C) Adult brains dissected from *elav*>GFP-caz (A), *elav*>GFP-Smn (B) and *elav*>GFP-TBPH (C) flies 5-7 days after expression. The GFP signal is shown in green. Insets in a1-a3, b1-b3 and c1-c3 show the sub-cellular distribution of the GFP-tagged proteins. GFP signals are shown in white (left) or green (overlay, right). DAPI signals are shown in white (middle) or blue (overlay, right). Scale bar: 50 μm. Complete genotypes: *elav*-Gal4/Y; tub-Gal80ts/UAS-GFP-*caz* (A), *elav*-Gal4/Y; tub-Gal80ts/UAS-GFP-*Smn* (B) and *elav*-Gal4/Y; tub-Gal80ts/UAS-GFP-*TBPH* (C). (D) Western blot performed on lysates from adult flies with pan neural (*elav-*Gal4) expression of GFP-Caz (left), GFP-Smn (middle) and GFP-TBPH (right) brains. Anti-GFP antibodies were used to detect GFP fusions. Tubulin was used as a loading control. (E) qRT-PCR quantification of caz, Smn, and TBPH in VDRC strain #13673 expressing dsRNA targeting *always early* (contr.) and flies containing the shRNA transgene for each target gene and grown in the presence of hormone for 10 days. ddCt values to the endogenous control gene Rp49 were normalized to the corresponding ddCts from samples of flies grown in the absence of hormone. The black bar for the control samples represents the average normalized expression of the three target genes in the VDRC strain.

**FigS1.pdf**

### Additional file 2

**Supplementary Data 1.** Sequencing library statistics

**Supdata1.xlxs**

### Additional file 3

**Supplementary Figure S2. Overview of sequencing datasets.** Sample-to-sample distance heatmap for (A) the RIP-seq and (B) the RNA-Seq datasets, revealing overall similarities and dissimilarities between dataset samples based on Euclidean distance. (C) Principal component analysis for RNA-Seq datasets. Left: full dataset, samples colored by treatment, symbols indicate fly line (condition). Right: analysis of hormone treated samples, colored by fly line. (D) Volcano plots of DE genes identified for each knockdown (with adj. p value < 0.05). Genes displaying common changes across the three fly lines are highlighted in color. (E) Total number of DE genes identified in each sample type at increasing levels of | log2 FC |. (F) Number of common genes between the three knockdowns identified at increasing | log2 FC | cut-offs. The expected versus observed number of common genes in the overlaps is displayed in red and black, respectively. The percentage represented by the common genes in the smallest dataset of the overlap (Smn DE) is shown over the plot bars. Increasing fold change cut-offs leads to a progressively bigger reduction of “captured” genes without benefits to the functional enrichment analysis.

**FigS2.pdf**

### Additional file 4

**Supplementary Data 2.** List of RIP-Seq enriched transcripts

**Supdata2.xlxs**

### Additional file 5

**Supplementary Figure S3: Coverage of RIP-Seq and RNA-Seq experiments in FlyAtlas tissue-specific expression profiles.** (A) Normalized RNA-Seq data of adult fly brain tissue was retrieved from the FlyAtlas2 database (see methods). The total 9020 transcripts were filtered using an expression threshold of > 1 FPKM. From the total 7369 transcripts identified in the RIP-Seq and knockdown experiments, 5511 were also detected in this dataset, and will be referred to as “neuronal” transcripts hereafter. Bar graph shows the number of transcripts identified in each experiment. (B) Figure summarizes the top 7 functions enriched in the sets of neuronal and non-neuronal transcripts identified in RIP-Seq and knockdown experiments. clusterProfiler R package was used to compare the functional enrichment of the 5511 “neuronal” and 1858 “non-neuronal” transcripts using Gene Ontology Biological Process, hyper-geometric test, adjusted p-value 0.05. From 824 enriched terms in neuronal transcripts, 92 include at the description the following key words: “synap”, “axon”, “neuro”, “dendrite”, “nervous”, “button”, “glial” or “cortex”. Non-neuronal transcripts were enriched in 19 terms, none of them related to neuronal processes. (C) Figure shows density plots of log_2_-transformed FPKM values for transcripts identified in the RIP-Seq and RNA-Seq experiments and classified as “non-neuronal”. 67.4% of the 1858 transcripts were detected in 10 additional tissues available in the FlyAtlas2 and displayed highest expression densities on *head*, *thoracicoabdominal ganglion* and *eye* tissues, explaining their possible origin in the datasets. (D) Density plot of log_2_-transformed FPKM values of “neuronal” transcripts from the FlyAtlas2, RIP-Seq, DE/AS, and selected functional modules subsets, revealing an enrichment of our datasets in transcripts with medium to high expression levels in neurons, particularly for the transcripts with altered expression retained in the selected modules.

**FigS3.pdf**

### Additional file 6

**Supplementary Data 3.** RNA-Seq DE transcripts

**Supdata3.xlxs**

### Additional file 7

**Supplementary Data 4.** Alternative splicing analysis results

**Supdata4.xlxs**

### Additional file 8

**Supplementary Data 5.** Annotation of all FlyAtlas “neuronal” genes regarding the presence in the different DE/AS/RIP data subsets

**Supdata5.xlxs**

### Additional file 9

**Supplementary Data 6.** Functional Enrichment Analysis of target gene-dependent transcripts

**Supdata6.xlxs**

### Additional file 10

**Supplementary Data 7.** Functional Module annotation

**Supdata7.xlxs**

### Additional file 11

**Supplementary Figure S4: Evaluation of protein redundancy across functional modules.** (A) Complete-linkage hierarchical clustering using Jaccard’s similarity coefficient for the 122 modules with a size between 10 to 100 proteins. The 52 modules passing the overall impact cut-off of >20% of transcripts altered in at least one knockdown are labeled in red. (B) Box plots describing the number of modules sharing at least one protein when comparing modules including less or more than 100 proteins Wilcoxon test, p value 2.2×10^-16^. (C) Bar plot indicating the number of proteins found in common between different super-modules. Colored horizontal bars indicate total number of proteins in each super-module. Black vertical bars indicate the overlap between super-modules. Only overlap sets including at least 5 proteins are shown.

### Additional file 12

**Supplementary Data 8.** NMJ Module

**Supdata8.xlxs**

### Additional file 13

**Supplementary Methods.** Primers used for cloning and qRT-PCR

**SupMethods.pdf**

## References

Achsel T, Barabino S, Cozzolino M, Carrì MT. 2013. The intriguing case of motor neuron disease: ALS and SMA come closer. Biochemical Society transactions 41: 1593–1597.

Alonso-López D., Gutiérrez M.A., Lopes K.P. et al. 2016. APID interactomes: providing proteome-based interactomes with controlled quality for multiple species and derived networks. Nucleic Acids Research 44(W1): W529–W535.

Alonso-López D, Campos-Laborie FJ, Gutiérrez MA, Lambourne L, Calderwood MA, Vidal M, De Las Rivas J. 2019. APID database: redefining protein-protein interaction experimental evidences and binary interactomes. Database (Oxford) 2019: baz005.

Amaral AJ, Brito FF, Chobanyan T, Yoshikawa S, Yokokura T, Van Vactor D, Gama-Carvalho M. 2014. Quality assessment and control of tissue specific RNA-seq libraries of Drosophila transgenic RNAi models. Frontiers in genetics 5: 43.

Aquilina B, Cauchi RJ. 2018. Modelling motor neuron disease in fruit flies: Lessons from spinal muscular atrophy. Journal of neuroscience methods 310: 3–11.

Birsa N, Bentham MP, Fratta P. 2020. Cytoplasmic functions of TDP-43 and FUS and their role in ALS. Seminars in cell & developmental biology 99: 193–201.

Boulisfane N, Choleza M, Rage F, Neel H, Soret J, Bordonné R. 2011. Impaired minor tri-snRNP assembly generates differential splicing defects of U12-type introns in lymphoblasts derived from a type I SMA patient. Human molecular genetics 20: 641–648.

Bowerman M, Murray LM, Scamps F, Schneider BL, Kothary R, Raoul C. 2018. Pathogenic commonalities between spinal muscular atrophy and amyotrophic lateral sclerosis: Converging roads to therapeutic development. European journal of medical genetics 61: 685–698.

Cacciottolo R, Ciantar J, Lanfranco M, Borg RM, Vassallo N, Bordonné R, Cauchi RJ. 2019. SMN complex member Gemin3 self-interacts and has a functional relationship with ALS-linked proteins TDP-43, FUS and Sod1. Scientific reports 9: 18666.

Chang HC, Dimlich DN, Yokokura T, Mukherjee A, Kankel MW, Sen A, Sridhar V, Fulga TA, Hart AC, Van Vactor D et al. 2008. Modeling spinal muscular atrophy in Drosophila. PloS one 3: e3209.

Chi B, O’Connell JD, Iocolano AD, Coady JA, Yu Y, Gangopadhyay J, Gygi SP, Reed R. 2018. The neurodegenerative diseases ALS and SMA are linked at the molecular level via the ASC-1 complex. Nucleic acids research 46: 11939–11951.

Cragnaz L, Klima R, Skoko N, Budini M, Feiguin F, Baralle FE. 2014. Aggregate formation prevents dTDP-43 neurotoxicity in the Drosophila melanogaster eye. Neurobiology of disease 71: 74–80.

Da Cruz S, Cleveland DW. 2011. Understanding the role of TDP-43 and FUS/TLS in ALS and beyond. Current opinion in neurobiology 21: 904–919.

Dobin A, Davis CA, Schlesinger F, Drenkow J, Zaleski C, Jha S, Batut P, Chaisson M, Gingeras TR. 2013. STAR: ultrafast universal RNA-seq aligner. Bioinformatics (Oxford, England) 29: 15–21.

Donlin-Asp PG, Bassell GJ, Rossoll W. 2016. A role for the survival of motor neuron protein in mRNP assembly and transport. Current opinion in neurobiology 39: 53–61.

Donlin-Asp PG, Fallini C, Campos J, Chou CC, Merritt ME, Phan HC, Bassell GJ, Rossoll W. 2017. The Survival of Motor Neuron Protein Acts as a Molecular Chaperone for mRNP Assembly. Cell reports 18: 1660–1673.

dos Santos G, Schroeder AJ, Goodman JL, Strelets VB, Crosby MA, Thurmond J, Emmert DB, Gelbart WM. 2015. FlyBase: introduction of the Drosophila melanogaster Release 6 reference genome assembly and large-scale migration of genome annotations. Nucleic acids research 43: D690–697.

Ederle H, Dormann D. 2017. TDP-43 and FUS en route from the nucleus to the cytoplasm. FEBS letters 591: 1489–1507.

Fallini C, Donlin-Asp PG, Rouanet JP, Bassell GJ, Rossoll W. 2016. Deficiency of the Survival of Motor Neuron Protein Impairs mRNA Localization and Local Translation in the Growth Cone of Motor Neurons. The Journal of neuroscience : the official journal of the Society for Neuroscience 36: 3811–3820.

Fallini C, Zhang H, Su Y, Silani V, Singer RH, Rossoll W, Bassell GJ. 2011. The survival of motor neuron (SMN) protein interacts with the mRNA-binding protein HuD and regulates localization of poly(A) mRNA in primary motor neuron axons. The Journal of neuroscience : the official journal of the Society for Neuroscience 31: 3914–3925.

Fernandes N, Eshleman N, Buchan JR. 2018. Stress Granules and ALS: A Case of Causation or Correlation? Advances in neurobiology 20: 173–212.

Gama-Carvalho M, Garcia-Vaquero M, Pinto FR, Besse F, Weis J, Voigt A, Schulz JB, De Las Rivas J. 2017. Linking amyotrophic lateral sclerosis and spinal muscular atrophy through RNA-transcriptome homeostasis: a genomics perspective. Journal of neurochemistry 141: 12–30.

Garcia-Vaquero M, Gama-Carvalho M, Pinto FR, De Las Rivas J. 2022. Biological Interacting Units identified in human protein networks reveal tissue functional diversification and its impact on disease. Computational and Structural Biotechnology Journal 20: 3764–3778.

Grice SJ, Liu JL. 2011. Survival motor neuron protein regulates stem cell division, proliferation, and differentiation in Drosophila. PLoS genetics 7: e1002030.

Groen EJ, Fumoto K, Blokhuis AM, Engelen-Lee J, Zhou Y, van den Heuvel DM, Koppers M, van Diggelen F, van Heest J, Demmers JA, et al. 2013. ALS-associated mutations in FUS disrupt the axonal distribution and function of SMN. Human molecular genetics 22: 3690–3704.

Huber W, Carey VJ, Gentleman R, Anders S, Carlson M, Carvalho BS, Bravo HC, Davis S, Gatto L, Girke T et al. 2015. Orchestrating high-throughput genomic analysis with Bioconductor. Nature methods 12: 115–121.

Kankel MW, Sen A, Lu L, Theodorou M, Dimlich DN, McCampbell A, Henderson CE, Shneider NA, Artavanis-Tsakonas S. 2020. Amyotrophic Lateral Sclerosis Modifiers in Drosophila Reveal the Phospholipase D Pathway as a Potential Therapeutic Target. Genetics 215: 747–766.

Kline RA, Kaifer KA, Osman EY, Carella F, Tiberi A, Ross J, Pennetta G, Lorson CL, Murray LM. 2017. Comparison of independent screens on differentially vulnerable motor neurons reveals alpha-synuclein as a common modifier in motor neuron diseases. PLoS genetics 13: e1006680.

Lagier-Tourenne C, Polymenidou M, Hutt KR, Vu AQ, Baughn M, Huelga SC, Clutario KM, Ling SC, Liang TY, Mazur C et al. 2012. Divergent roles of ALS-linked proteins FUS/TLS and TDP-43 intersect in processing long pre-mRNAs. Nature neuroscience 15: 1488–1497.

Law CW, Chen Y, Shi W, Smyth GK. 2014. voom: Precision weights unlock linear model analysis tools for RNA-seq read counts. Genome biology 15: R29.

Leader DP, Krause SA, Pandit A, Davies SA, Dow JAT. 2018. FlyAtlas 2: a new version of the Drosophila melanogaster expression atlas with RNA-Seq, miRNA-Seq and sex-specific data. Nucleic acids research 46: D809–d815.

Li DK, Tisdale S, Lotti F, Pellizzoni L. 2014. SMN control of RNP assembly: from post-transcriptional gene regulation to motor neuron disease. Seminars in cell & developmental biology 32: 22–29.

Li YR, King OD, Shorter J, Gitler AD. 2013. Stress granules as crucibles of ALS pathogenesis. The Journal of cell biology 201: 361–372.

Liguori F, Amadio S, Volonté C. 2021. Fly for ALS: Drosophila modeling on the route to amyotrophic lateral sclerosis modifiers. Cellular and molecular life sciences : CMLS 78: 6143–6160.

Ling SC, Polymenidou M, Cleveland DW. 2013. Converging mechanisms in ALS and FTD: disrupted RNA and protein homeostasis. Neuron 79: 416–438.

Love MI, Huber W, Anders S. 2014. Moderated estimation of fold change and dispersion for RNA-seq data with DESeq2. Genome biology 15: 550.

McGuire SE, Le PT, Osborn AJ, Matsumoto K, Davis RL. 2003. Spatiotemporal rescue of memory dysfunction in Drosophila. Science (New York, NY) 302: 1765–1768.

McGurk L, Berson A, Bonini NM. 2015. Drosophila as an In Vivo Model for Human Neurodegenerative Disease. Genetics 201: 377–402.

Mili S, Steitz JA. 2004. Evidence for reassociation of RNA-binding proteins after cell lysis: implications for the interpretation of immunoprecipitation analyses. RNA (New York, NY) 10: 1692–1694.

Mora A, Donaldson IM. 2011. iRefR: an R package to manipulate the iRefIndex consolidated protein interaction database. BMC bioinformatics 12: 455.

Olesnicky EC, Wright EG. 2018. Drosophila as a Model for Assessing the Function of RNA-Binding Proteins during Neurogenesis and Neurological Disease. Journal of developmental biology 6.

Osterwalder T, Yoon KS, White BH, Keshishian H. 2001. A conditional tissue-specific transgene expression system using inducible GAL4. Proceedings of the National Academy of Sciences of the United States of America 98: 12596–12601.

Pellizzoni L, Charroux B, Rappsilber J, Mann M, Dreyfuss G. 2001. A functional interaction between the survival motor neuron complex and RNA polymerase II. The Journal of cell biology 152: 75–85.

Perera ND, Sheean RK, Crouch PJ, White AR, Horne MK, Turner BJ. 2016. Enhancing survival motor neuron expression extends lifespan and attenuates neurodegeneration in mutant TDP-43 mice. Human molecular genetics 25: 4080–4093.

Price PL, Morderer D, Rossoll W. 2018. RNP Assembly Defects in Spinal Muscular Atrophy. Advances in neurobiology 20: 143–171.

Ratti A, Buratti E. 2016. Physiological functions and pathobiology of TDP-43 and FUS/TLS proteins. Journal of neurochemistry 138 Suppl 1: 95–111.

Ritchie ME, Phipson B, Wu D, Hu Y, Law CW, Shi W, Smyth GK. 2015. limma powers differential expression analyses for RNA-sequencing and microarray studies. Nucleic acids research 43: e47.

Scialo F, Sriram A, Stefanatos R, Sanz A. 2016. Practical Recommendations for the Use of the GeneSwitch Gal4 System to Knock-Down Genes in Drosophila melanogaster. PloS one 11: e0161817.

Sen A, Dimlich DN, Guruharsha KG, Kankel MW, Hori K, Yokokura T, Brachat S, Richardson D, Loureiro J, Sivasankaran R, Curtis D, Davidow LS, Rubin LL, Hart AC, Van Vactor D, Artavanis-Tsakonas S. 2013. Genetic circuitry of Survival motor neuron, the gene underlying spinal muscular atrophy. Proceedings of the National Academy of Sciences of the United States of America 110: E2371–80.

Shen S, Park JW, Lu ZX, Lin L, Henry MD, Wu YN, Zhou Q, Xing Y. 2014. rMATS: robust and flexible detection of differential alternative splicing from replicate RNA-Seq data. Proceedings of the National Academy of Sciences of the United States of America 111: E5593–5601.

Spradling AC, Stern D, Beaton A, Rhem EJ, Laverty T, Mozden N, Misra S, Rubin GM. 1999. The Berkeley Drosophila Genome Project gene disruption project: Single P-element insertions mutating 25% of vital Drosophila genes. Genetics 153: 135–177.

Spring AM, Raimer AC, Hamilton CD, Schillinger MJ, Matera AG. 2019. Comprehensive Modeling of Spinal Muscular Atrophy in Drosophila melanogaster. Frontiers in molecular neuroscience 12: 113.

Sun S, Ling SC, Qiu J, Albuquerque CP, Zhou Y, Tokunaga S, Li H, Qiu H, Bui A, Yeo GW et al. 2015. ALS-causative mutations in FUS/TLS confer gain and loss of function by altered association with SMN and U1-snRNP. Nature communications 6: 6171.

Taylor JP, Brown RH, Jr., Cleveland DW. 2016. Decoding ALS: from genes to mechanism. Nature 539: 197–206.

Tisdale S, Lotti F, Saieva L, Van Meerbeke JP, Crawford TO, Sumner CJ, Mentis GZ, Pellizzoni L. 2013. SMN is essential for the biogenesis of U7 small nuclear ribonucleoprotein and 3’-end formation of histone mRNAs. Cell reports 5: 1187–1195.

Tsuiji H, Iguchi Y, Furuya A, Kataoka A, Hatsuta H, Atsuta N, Tanaka F, Hashizume Y, Akatsu H, Murayama S et al. 2013. Spliceosome integrity is defective in the motor neuron diseases ALS and SMA. EMBO molecular medicine 5: 221–234.

Vijayakumar J, Perrois C, Heim M, Bousset L, Alberti S, Besse F. 2019. The prion-like domain of Drosophila Imp promotes axonal transport of RNP granules in vivo. Nature communications 10: 2593.

Wang JW, Brent JR, Tomlinson A, Shneider NA, McCabe BD. 2011. The ALS-associated proteins FUS and TDP-43 function together to affect Drosophila locomotion and life span. The Journal of clinical investigation 121: 4118–4126.

Wang M, Chen PY, Wang CH, Lai TT, Tsai PI, Cheng YJ, Kao HH, Chien CT. 2016. Dbo/Henji Modulates Synaptic dPAK to Gate Glutamate Receptor Abundance and Postsynaptic Response. PLoS genetics 12: e1006362.

Wang M, Zhao Y, Zhang B. 2015. Efficient Test and Visualization of Multi-Set Intersections. Scientific reports 5: 16923.

Wickham H. 2016. ggplot2: Elegant Graphics for Data Analysis. Springer-Verlag New York. ISBN 978-3-319-24277-4, https://ggplot2.tidyverse.org

Workman E, Kolb SJ, Battle DJ. 2012. Spliceosomal small nuclear ribonucleoprotein biogenesis defects and motor neuron selectivity in spinal muscular atrophy. Brain research 1462: 93–99.

Xia R, Liu Y, Yang L, Gal J, Zhu H, Jia J. 2012. Motor neuron apoptosis and neuromuscular junction perturbation are prominent features in a Drosophila model of Fus-mediated ALS. Molecular neurodegeneration 7: 10.

Yamazaki T, Chen S, Yu Y, Yan B, Haertlein TC, Carrasco MA, Tapia JC, Zhai B, Das R, Lalancette-Hebert M et al. 2012. FUS-SMN protein interactions link the motor neuron diseases ALS and SMA. Cell reports 2: 799–806.

Zaepfel BL, Rothstein JD. 2021. RNA Is a Double-Edged Sword in ALS Pathogenesis. Frontiers in cellular neuroscience 15: 708181.

Zbinden A, Pérez-Berlanga M, De Rossi P, Polymenidou M. 2020. Phase Separation and Neurodegenerative Diseases: A Disturbance in the Force. Developmental cell 55: 45–68.

Zhang Z, Lotti F, Dittmar K, Younis I, Wan L, Kasim M, Dreyfuss G. 2008. SMN deficiency causes tissue-specific perturbations in the repertoire of snRNAs and widespread defects in splicing. Cell 133: 585–600.

Zhao DY, Gish G, Braunschweig U, Li Y, Ni Z, Schmitges FW, Zhong G, Liu K, Li W, Moffat J et al. 2016. SMN and symmetric arginine dimethylation of RNA polymerase II C-terminal domain control termination. Nature 529: 48–53.

Zou J, Barahmand-pour F, Blackburn ML, Matsui Y, Chansky HA, Yang L. 2004. Survival motor neuron (SMN) protein interacts with transcription corepressor mSin3A. The Journal of biological chemistry 279: 14922–14928.

