## Supplementary Methods for "Analysis of pre-symptomatic *Drosophila* models for ALS and SMA reveals convergent impact on functional protein complexes linked to neuro-muscular degeneration"

**Supplementary Methods** – Primer sequences

| **Primer name** | **Sequence (5’-3’)** |
|---|---|
| Smn_fwd_cloning | CACCATGTCCGACGAGACGAACG |
| Smn_rev_cloning | GATGGAATTACTTCTTGGGTGTC |
| Caz fwd_cloning | CACCATGGAACGTGGCGGTTATGGTG |
| Caz_rev_cloning | TTAATATGGTCTCGAGCGCATGC |
| NotI_TBPH-fwd | AAAAGCGGCCGCCATGGATTTCGTTCAAG |
| XhoI_TBPH_rev | AAAACTCGAGTTAAAGAAAGTTTGACTTCTCCGC |
| RT-PCR caz_rev | TCCGCGATCGAAGCGACCTCC |
| RT-PCR caz_for | TCCTACGGAAATGGAGGCGCC |
| RT-PCR TBPH_for | GGAAGGGGCCGAATAACCCGAAC |
| RT-PCR TBPH_rev | CACACATCATTGGGTGACAGGCACC |
| RT-PCR Smn1_ for | AAGAAGAATGCCACAACTCCC |
| RT-PCR Smn1_ rev | CAATGGACGTAATAGTAGCTGGG |
| RT-PCR rp49_for | TCGGATCGATATGCTAAGCTGTCGCAC |
| RT-PCR rp49_rev | AGGCGACCGTTGGGGTTGGTGAG |
| RT-PCR act5C_for | CACACCGTGCCCATCTACGAG |
| RT-PCR act5C_rev | CTTCTGCATACGGTCGGCGATGC |
