## Supplementary figures and images for "Analysis of pre-symptomatic *Drosophila* models for ALS and SMA reveals convergent impact on functional protein complexes linked to neuro-muscular degeneration"

### Supplementary Fig. S1

Supplementary Figure 1

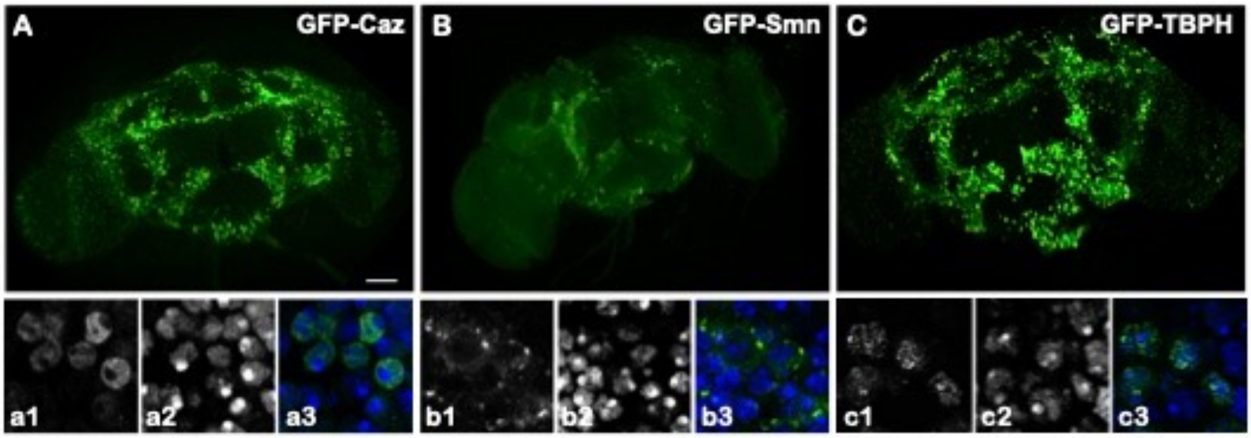


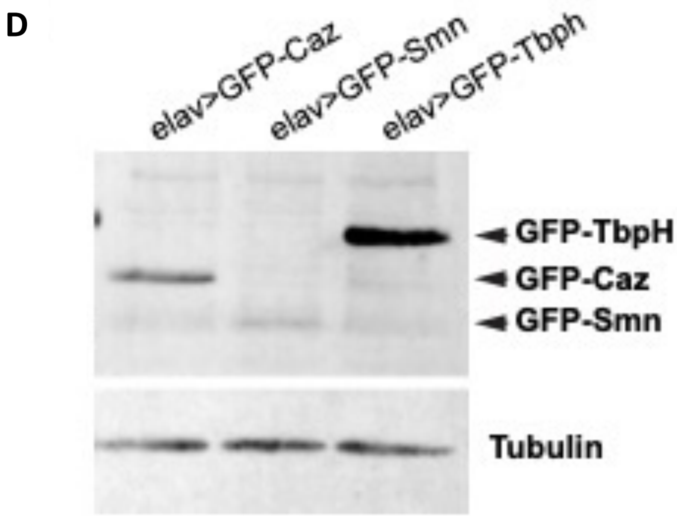


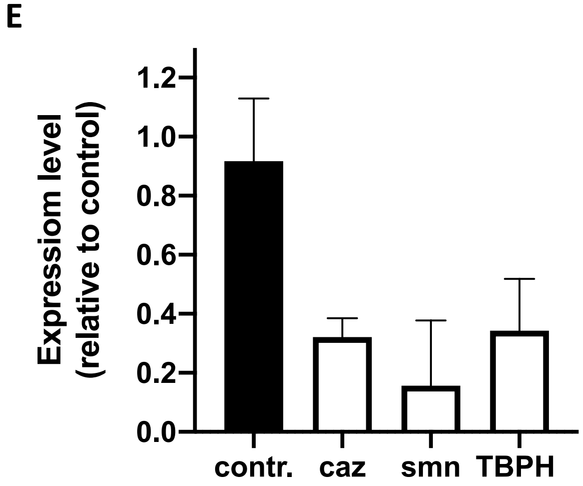

### Supplementary Fig. S2

#### Supplementary Figure 2

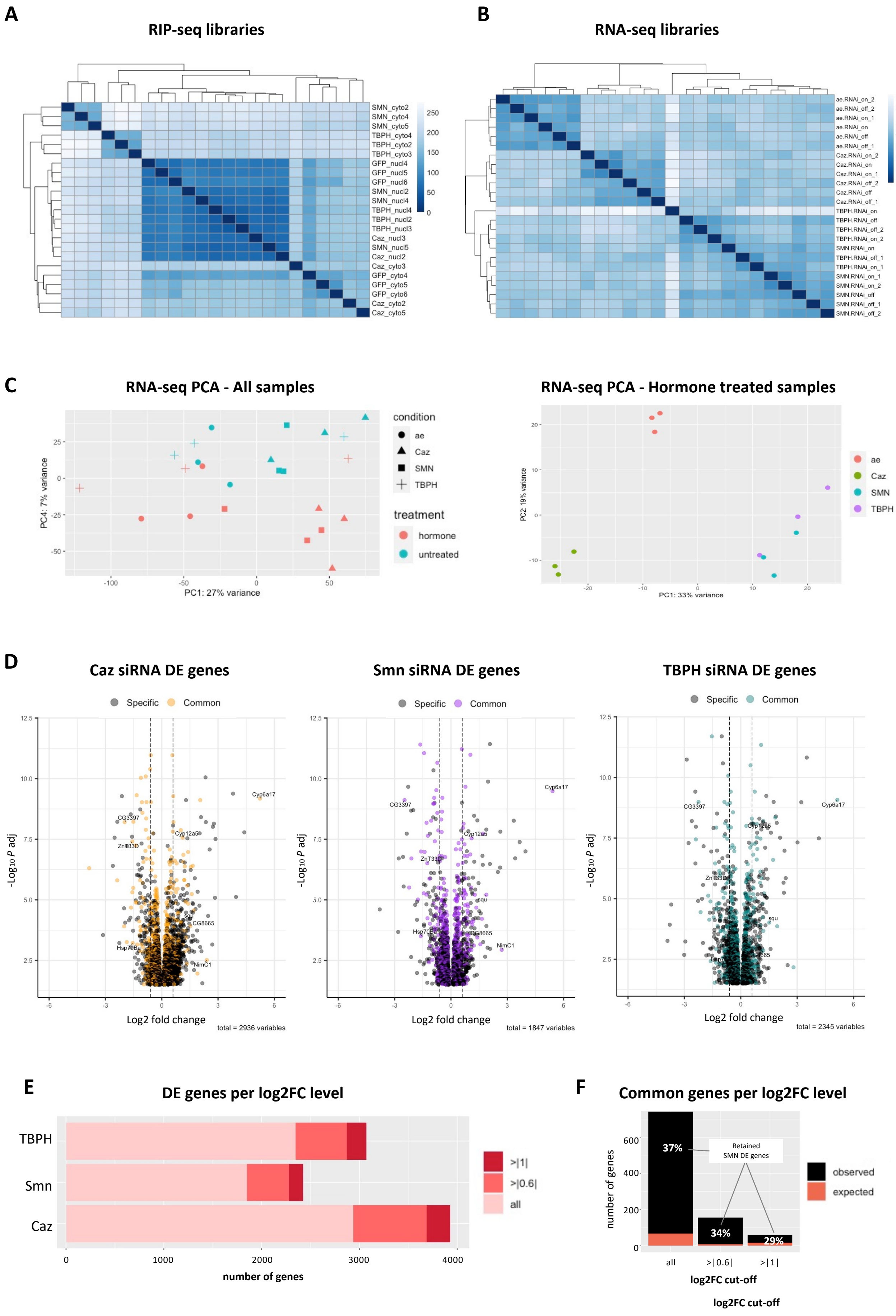

### Supplementary Fig. S3

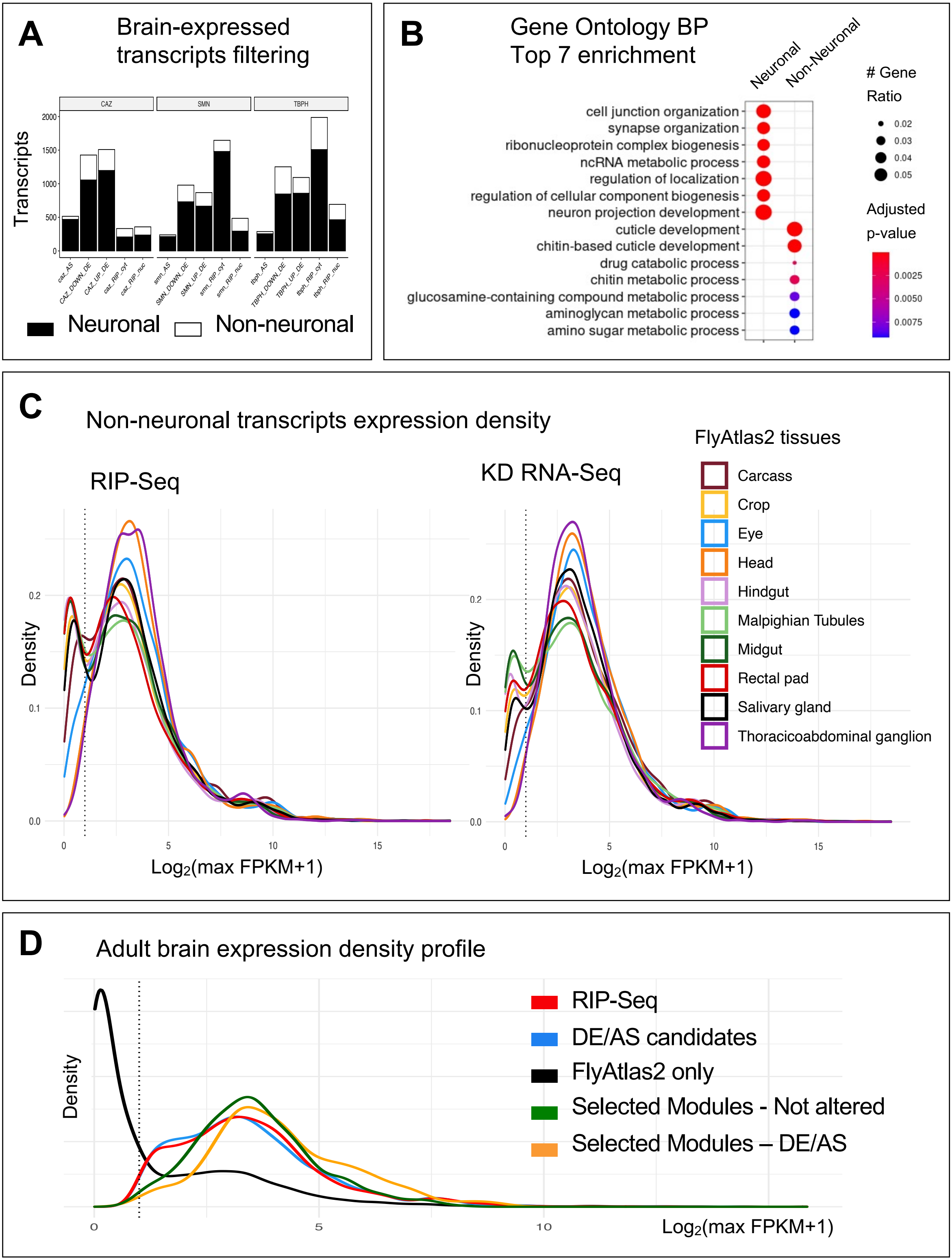

### Supplementary Fig. S4

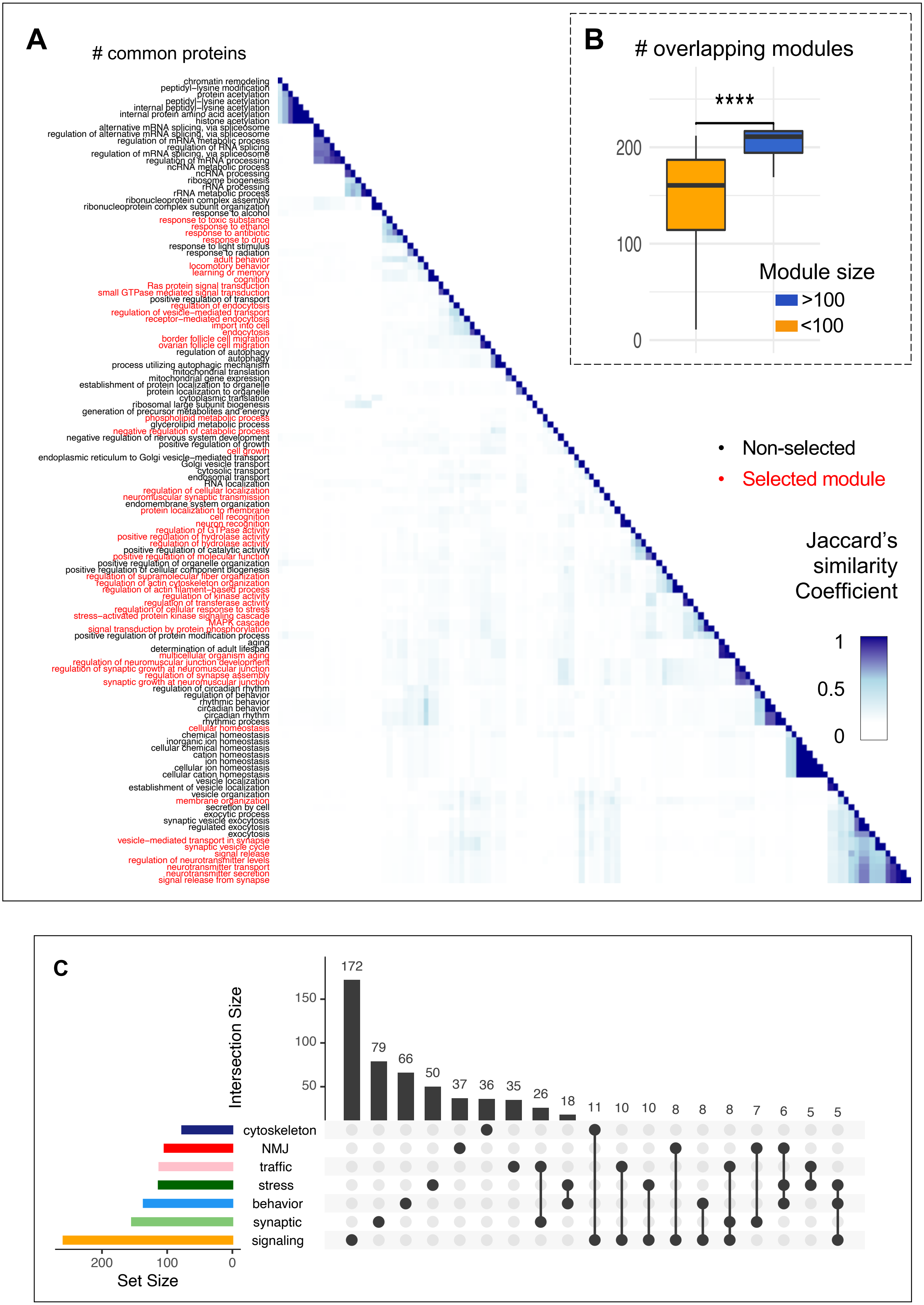
